# Noradrenergic-dependent restoration of visual discrimination in a mouse model of *SYNGAP1*-related disorder

**DOI:** 10.1101/2025.04.09.647923

**Authors:** Danai Katsanevaki, Nathalie Dupuy, Sam A Booker, Damien Wright, Aisling Kenny, Zihao Chen, Nina Kudryashova, Pippa Howitt, Andrew C Stanfield, Peter C Kind, Nathalie L Rochefort

## Abstract

Atypical sensory processing in neurodevelopmental disorders contributes to cognitive, social, and behavioural disruptions, yet underlying neurophysiological mechanisms remain unclear. Using a mouse model of *SYNGAP1* haploinsufficiency (HET), a common monogenic cause of intellectual disability and autism, we investigated visual processing deficits. *Syngap* HET mice exhibited impaired behavioural visual discriminability, associated with reduced coding precision for visual stimuli in the primary visual cortex (V1). Notably, intrinsic properties of V1 neurons and visual responses under anaesthesia were unaltered, suggesting behavioural state-dependent disruptions in awake *Syngap* HET mice. Supporting this, both mice and individuals with *SYNGAP1* haploinsufficiency exhibited larger pupil size during visual stimulation, implicating neuromodulatory dysfunction. Targeting noradrenergic tone systemically with an α_2_-adrenergic receptor agonist restored V1 coding precision in *Syngap* HET mice. Our findings reveal neuromodulatory dysregulation as a novel mechanism underlying sensory disruptions in *SYNGAP1*-related disorder, highlighting potential therapeutic targets for addressing sensory impairments in neurodevelopmental disorders.

## Introduction

Atypical sensory processing is increasingly recognised as a key feature in syndromic and idiopathic forms of neurodevelopmental disorders, including autism spectrum disorders (ASD)^1^. These sensory disruptions lead to, and often predict, a range of cognitive, social, and behavioural affective impairments^2–4^, strongly impacting the quality of life of affected individuals. However, despite the prevalence and importance of sensory processing abnormalities in neurodevelopmental disorders^5^, the underlying neurophysiological mechanisms that disrupt sensory perception remain widely understudied.

Large-scale exome sequencing studies have identified *SYNGAP1* as one of the most prevalent neurodevelopmental disorder genes, accounting for up to 1% of all affected individuals^6–9^. *SYNGAP1* encodes a synaptically enriched protein, SYNGAP, that is essential for survival and development^10–15^. Individuals with *de novo* pathogenic mutations in *SYNGAP1* experience global developmental delay, with moderate-to-severe intellectual disability^9,16–18^, and often meet the diagnostic criteria for ASD^17,19–23^. Nearly all individuals with *SYNGAP1*-related disorder exhibit atypical sensory processing^24^, spanning multiple sensory modalities including tactile, auditory, proprioceptive, gustatory and visual perception^23,25–30^. Moreover, individuals with *SYNGAP1*-related disorder frequently display atypically high scores in all four quadrants measured by a Short Sensory Profile-2 (SSP-2)^31^, namely seeking, avoiding, sensitivity, and registration^32^.

A powerful approach to explore the neuronal mechanisms underlying these sensory and cognitive disruptions is to use animal models of single-gene mutations^26,33–39^. In this study, we used a mouse model of *SYNGAP1* haploinsufficiency to investigate the mechanisms underlying visual disruptions. Using two-photon calcium imaging in the primary visual cortex (V1) of adult mice, we found that V1 neurons of heterozygous *Syngap* (HET) mice exhibited greater variability in response to visual stimuli, leading to reduced coding precision compared to wild-type (WT) littermate controls. These cortical deficits were associated with reduced behavioural visual discriminability. *In vitro* recordings showed that the intrinsic or synaptic properties of V1 neurons were not affected in *Syngap* HET mice. In addition, V1 visual responses under anaesthesia were found to be similar in both WT and *Syngap* HET mice. These results, combined with the observation of differences in pupil diameter, suggested a dysregulation of behavioural state modulation in *Syngap* HET mice. Since behavioural state regulation strongly depends on neuromodulatory inputs, we tested the impact of targeting noradrenergic tone via systemic administration of an α_2_-adrenergic receptor agonist, guanfacine; this restored coding precision of visual stimuli in *Syngap* HET mice. Our findings reveal that dysregulations in behavioural state underlie visual deficits in *Syngap* HET mice and identify noradrenergic inputs as a novel mechanism underlying sensory processing deficits in *SYNGAP1*-related disorder.

## Results

In order to investigate the neuronal mechanisms underlying altered visual information processing in *SYNGAP1*-related disorder, we monitored the activity of layer 2/3 (L2/3) GCaMP6f-expressing neurons using *in vivo* two photon calcium imaging in the primary visual cortex (V1) of awake head-fixed adult *Syngap* heterozygous (HET) mice and wild-type (WT) littermate controls (Fig **1A** and see **Methods**).

**Figure 1.**
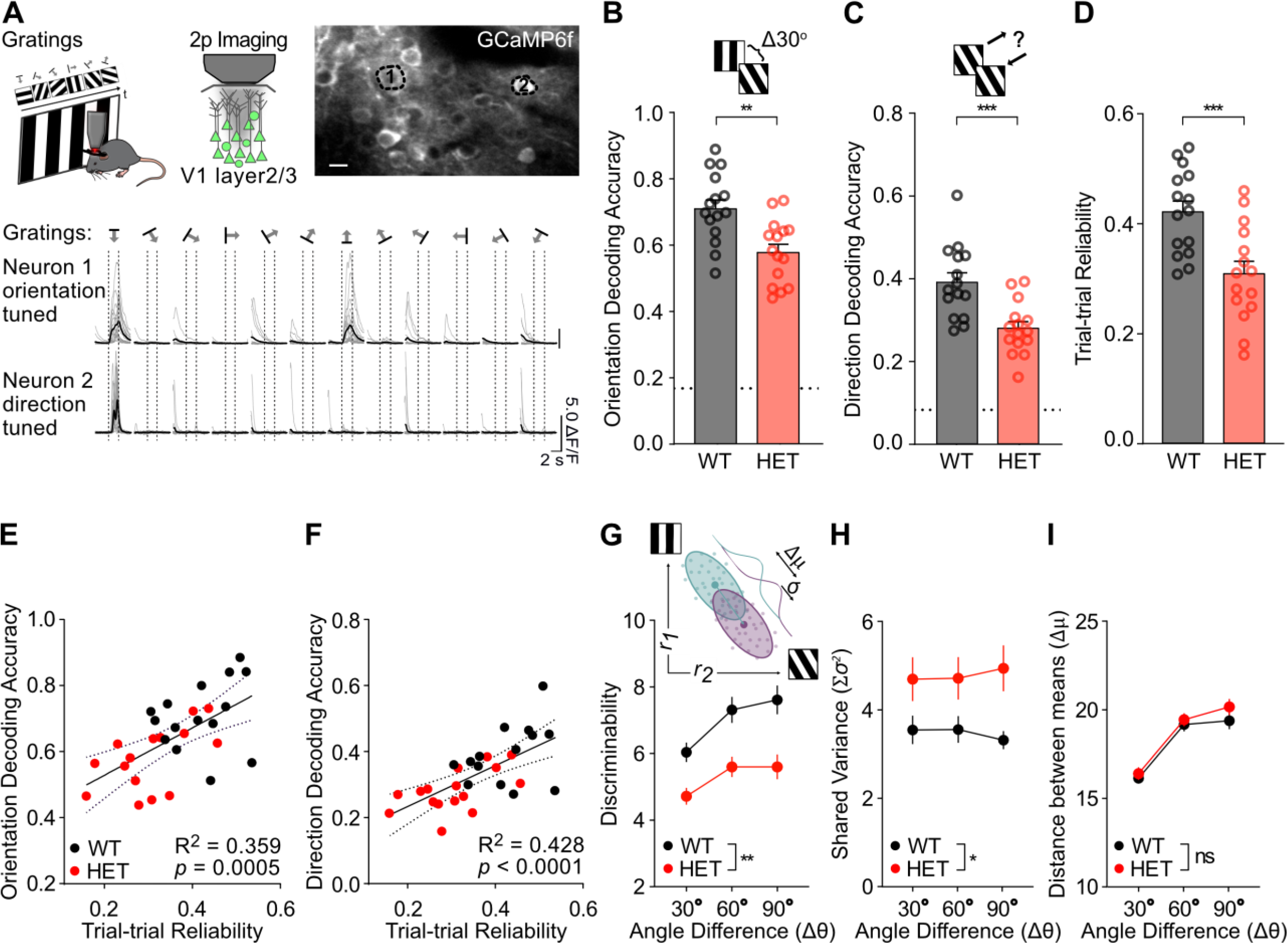
Reduced population coding precision and reduced reliability of visual responses in the primary visual cortex of *Syngap* HET mice. **(A)** (top) Schema of experimental procedure. Mice were head-fixed in front of a computer screen displaying drifting gratings, while GCamp6f-expressing layer 2/3 neurons were imaged in the primary visual cortex (V1). Example of *in vivo* field of view of GCaMP6f-labeled layer 2/3 neurons (scale bar: 10μm). (bottom) Example traces of GCaMP6f fluorescence changes over time (ΔF/F_0_) of an orientation-selective neuron (1) and a direction-selective neuron (2) (as shown above) during the presentation of 12 drifting gratings. Arrows indicate the angle of drift direction. Individual trials in grey; average of 20 trials in black. Scale bars: 2 s, 5 ΔF/F_0_. **(B)** Mean accuracy in decoding the correct orientation amongst 6 equally-spaced orientations (0-150°) based on layer 2/3 neuronal activity (maximum likelihood decoder; *p*=0.0013, *two-tailed unpaired t-test*; n = 15 WT and n = 15 *Syngap* HET mice). Dotted line: chance level. **(C)** Mean accuracy in decoding the correct direction amongst 12 directions (maximum likelihood decoder; *p*=0.0005, *two-tailed unpaired t-test*; n = 15 WT and n = 15 *Syngap* HET mice). Dotted line: chance level. **(D)** Mean trial-to-trial reliability of visual responses, (*p*=0.0009, *two-tailed unpaired t-test*; n = 15 WT and n = 15 *Syngap* HET mice). **(E)** Scatterplot of trial-to-trial reliability versus orientation decoding accuracy (R squared = 0.3595, *p*=0.0005, *F-test*). Black line represents linear regression fit and dotted lines the 95 % confidence bands. **(F)** Scatterplot of trial-to-trial reliability versus direction decoding accuracy (R squared = 0.4276, *p*<0.0001, *F-test*). Black line represents linear regression fit and dotted lines the 95 % confidence bands. **(G)** Inset: Schema of calculation of stimulus discriminability. Coloured dots represent two hypothetical scatterplots of single trial responses of a pair of neurons (response neuron 1 (*r_1_*), response neuron 2 (*r_2_*)), to the presentation of two gratings with different angles. The two corresponding Gaussian distributions are shown with the distance between the means vector (Δμ) and Standard Deviation (σ). Values for discriminability are calculated for pairs of gratings (angle difference Δθ; 30°, 60°, 90°; WT vs HET *p*=0.0013, *RM 2-way ANOVA*; n = 15 WT and n = 15 *Syngap* HET mice). **(H)** Mean shared variance (Sσ^2^) calculated from the stimulus discriminability (WT vs HET *p*=0.0266, *RM 2-way ANOVA*; n = 15 WT and n = 15 *Syngap* HET mice). **(I)** Mean distance between the means (Δμ) (WT vs HET *p*=0.3838, *RM 2-way ANOVA*; n = 15 WT and n = 15 *Syngap* HET mice). The same number of neurons per animal was used to assess decoding and discriminability. Mean across animals ± SEM is shown. \**p*<0.05, \*\**p*<0.01, \*\*\**p*<0.001, \*\*\*\**p*<0.0001.

### Reduced V1 population coding precision and reduced reliability of visual responses in *Syngap* HET mice

We monitored visual responses of V1 L2/3 neurons in *Syngap* HET and WT mice (Fig **1A**) during the randomised presentation of square-wave drifting gratings of 6 orientations (each with 2 opposite directions). To assess the neuronal population coding precision of these stimuli, we used a maximum likelihood estimator^40^ to decode grating orientations and directions from the recorded neuronal activity. We found lower decoding accuracy for both stimulus orientation and direction in *Syngap* HET mice (Fig **1B-C**, Suppl Fig **1A-B**). Consistent with this result, neuronal visual responses displayed lower depth-of-modulation^41^ in *Syngap* HET mice compared to WT controls (Suppl Fig **1C**), indicating either reduced orientation tuning or reduced response reliability (or both). By analysing visual responses of individual V1 neurons, we found that L2/3 neurons of *Syngap* HET mice responded less reliably to repeated presentations of the same stimuli (Fig **1D**, Suppl Fig **1H**), while their average stimulus orientation and direction selectivity was not significantly different from WT controls (Suppl Fig **1D**-**G**). Across mice, we found a significant correlation between the trial-to-trial reliability of visual responses and the population decoding accuracy for both orientation (Fig **1E**) and direction (Fig **1F**) of drifting gratings: V1 neurons in *Syngap* HET mice displaying both reduced reliability of responses and decreased decoding accuracy.

This reduction in V1 coding precision was confirmed by assessing the discriminability^42,43^ of V1 neuronal population representations. Our results showed that population discriminability was significantly reduced in *Syngap* HET mice compared to WT mice, for pairs of gratings 30, 60, and 90 degrees apart (Fig **1G**). This reduction in discriminability was due to an increased shared variance in population responses (Σσ^2^) of *Syngap* HET V1 neurons (Fig **1H**), while population orientation tuning was unaltered compared to WT controls (Δμ; Fig **1I**).

In these experiments, head-fixed mice were either placed in a tube (stationary) or on a wheel, freely running. Notably, we found significant reductions in trial-to-trial reliability and coding precision in *Syngap* HET mice for both tube- and wheel-experimental conditions, compared to WT controls (Suppl Fig **2A-H**). On the wheel, mice of both genotypes spent most of their time stationary (Suppl Fig **2I**), with no significant differences in self-initiated locomotion velocity or velocity variability between groups (Suppl Fig **2J-K**). Furthermore, we quantified the effect of locomotion on neuronal visual responses using the locomotion modulation index (LMI) and found no significant difference between WT and *Syngap* HET mice (Suppl Fig **2L**). These results indicate that the deficits observed in *Syngap* HET mice are not due to differences in locomotion or its effects on neuronal responses. Since no significant interaction was found across both conditions (wheel and tube) and genotype, we merged the results from both conditions in Figure 1 (see **Supplementary Table 1**).

Altogether, our findings show a reduction of V1 population coding precision for visual stimuli in *Syngap* HET mice due to a decreased reliability of visual responses.

### Impaired behavioural visual discriminability *in Syngap* HET mice

We next examined whether reduced coding precision in V1 of *Syngap* HET mice impacted visual discrimination behaviourally. We used a previously designed forced-choice modified water-maze task^44,45^ in which mice are presented with two high-contrast gratings and learn to discriminate a target (90° oriented grating) from a non-target (45° oriented grating) to escape the water (Fig **2A**; see **Methods**). *Syngap* HET mice required significantly more training sessions in comparison to their WT littermates to learn to discriminate the target *vs* the non-target orientation (45° apart): while WT mice reached a performance level above 80% after an average of 11 days of training, *Syngap* HET mice required five additional days (Suppl Fig **3A**). At day 16, both WT and *Syngap* HET mice reached stable high behavioural performance (>80%), with no significant difference between genotypes (Suppl Fig **3A** and Fig **2B**; WT: 88.75% ± 2.63 vs HET: 87.86% ± 3.25).

**Figure 2.**
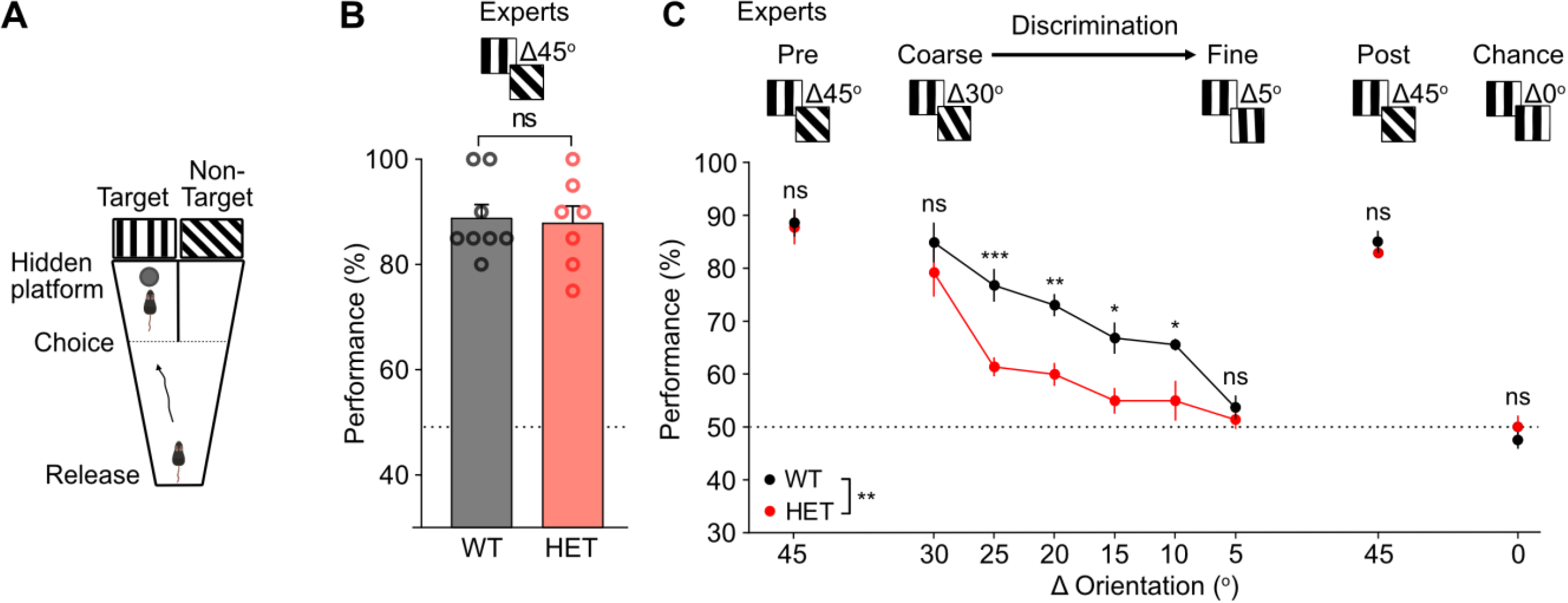
Impaired behavioural visual discriminability in *Syngap* HET mice. **(A)** Schema of behavioural task. Mice were placed in a modified water Y-maze and had to choose the arm associated with a target grating to reach a hidden platform and escape the water. **(B)** Task performance for 45° discrimination, at expert level. Mean performance of last two days of learning (*p*=0.8324, *two-tailed unpaired t-test*; n = 8 WT and n = 7 *Syngap* HET mice). Both genotypes learned the task and reached equal performance (see also Supplementary Fig. S3). **(C)** Testing of coarse to fine orientation discrimination. Behavioural performance as a function of discrimination difficulty, which was altered by changing the angle difference between target and non-target gratings (Δ Orientation; WT vs HET *p*=0.00531, *RM 2-way ANOVA*; n = 8 WT and n = 7 *Syngap* HET mice). Pre-Δ45° is the mean performance accuracy at the end of the training phase (same as panel **B**) and Post-Δ45° is a session done after the end of the testing. Dotted line: chance level. Mean across animals ± SEM is shown. \**p*<0.05, \*\**p*<0.01, \*\*\**p*<0.001, \*\*\*\**p*<0.0001.

Since after training both groups achieved expert performance for gratings 45° apart, we next tested fine visual discrimination by sequentially reducing the angle between the target and the non-target grating (Δ orientation). With increased difficulty, performance dropped for both groups, but the drop was more pronounced for *Syngap* HET mice that showed significantly lower performance for angle differences ≤ 25° compared to WT (Fig **2C**). Importantly, this was not due to a general decline in performance across successive days (*e.g.,* due to forgetting the task) since both groups still reached expert-level performance for the 45° angle discrimination at the end of the testing phase (WT: 85% ± 2.11 vs HET: 82.86% ± 1.01; Fig **2C** ’*45, post’*). Finally, differences in performance could not be explained by non-cognitive behavioural factors as assessed by the mean swimming speed during the task that was not different between both groups (Suppl Fig **3B-C**).

Collectively, these findings show that visual discriminability is impaired in *Syngap* HET compared to WT mice.

### Unaltered intrinsic and synaptic properties of V1 L2/3 pyramidal neurons of *Syngap* HET mice

We tested whether cell-intrinsic mechanisms, such as enhanced neuronal excitability, may account for the reduction of V1 population coding precision in *Syngap* HET mice. We used *in vitro* whole-cell patch clamp recordings from L2/3 neurons in V1 slices from adult mice (Fig **3A** **top**).

**Figure 3.**
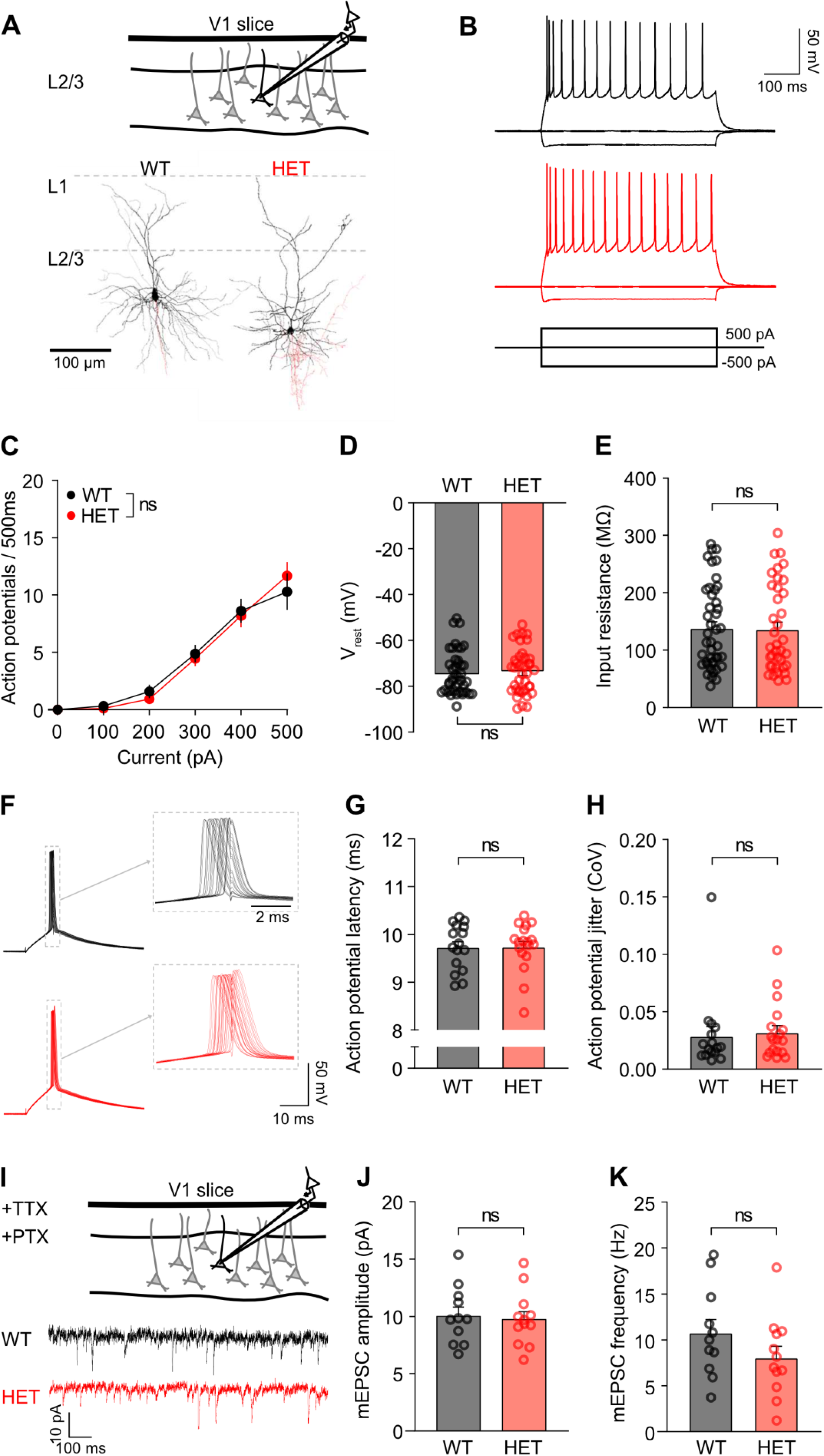
Unaltered intrinsic and synaptic properties of V1 L2/3 pyramidal neurons of *Syngap* HET mice *in vitro*. **(A)** (top) Schema of whole-cell patch-clamp recordings *in vitro*. (bottom) Flattened reconstructions of example V1 L2/3 pyramidal neurons. Somatodendritic (black) and axonal arborisation (red) are shown. Cortical layers are indicated by dashed grey lines. **(B)** Example firing characteristics of V1 L2/3 WT and *Syngap* HET neurons in response to 500 ms hyperpolarising to depolarising current injections (-500 to +500 pA, 100 pA steps). **(C)** Suprathreshold current-spike frequency (FI) responses of V1 L2/3 WT and *Syngap* HET neurons for current injections from 0-500 pA (WT vs HET *p*=0.893, *RM 2-way ANOVA*; n = 15 WT and n = 16 *Syngap* HET mice). **(D)** Mean resting membrane potential (*p*=0.5857, *Mann-Whitney test*; n = 43 WT cells from 15 mice and n = 39 *Syngap* HET cells from 16 mice). **(E)** Mean input resistance (*p*=0.8253, *Mann-Whitney test*; n = 43 WT cells from 15 mice and n = 39 *Syngap* HET cells from 16 mice). **(F)** Overlapped action potentials elicited at minimally suprathreshold currents (rheobase+25 pA, 10 ms) from WT (top) and *Syngap* HET (bottom) V1 L2/3 neurons. (inset) Expanded time-base for action potentials indicating the degree of jitter observed in both genotypes. **(G)** Mean action potential latency from the onset of the depolarising step (*p*=0.932, *Mann-Whitney test*; n = 16 WT cells from 7 mice and n = 18 *Syngap* HET cells from 7 mice). **(H)** Mean spike jitter (CoV; *p*=0.422, *Mann-Whitney test*; n = 16 WT cells from 7 mice and n = 18 *Syngap* HET cells from 7 mice). **(I)** (top) Schema of whole-cell patch-clamp recordings *in vitro*. (bottom) Example miniature EPSCs (mEPSCs) traces recorded in the presence of 300 nM TTX and 50 μM picrotoxin for a WT and a *Syngap* HET neuron. **(J)** Mean mEPSC amplitude (*p*=0.779, *two-tailed unpaired t-test*; n = 11 WT cells from 7 mice and n = 12 *Syngap* HET cells from 7 mice). **(K)** Mean mEPSC frequency (*p*=0.180, *two-tailed unpaired t-test*; n = 11 WT cells from 7 mice and n = 12 *Syngap* HET cells from 7 mice). Mean ± SEM is shown. \**p*<0.05, \*\**p*<0.01, \*\*\**p*<0.001, \*\*\*\**p*<0.0001.

Pyramidal cells were identified based on their responses to depolarising stimuli, and, in a subset of cells, morphological reconstructions (Fig **3A** **bottom**). We observed no significant differences in dendritic length or Sholl distribution between genotypes (**Suppl Fig 4A-F**), indicating that morphology of V1 L2/3 pyramidal neurons was unaltered in *Syngap* HET mice.

Similarly, intrinsic physiological properties of V1 L2/3 pyramidal cells were found to be unaltered in *Syngap* HET mice. Both responses to hyperpolarizing stimuli (Fig **3B**) and trains of repetitive action potentials in response to depolarizing stimuli showed no significant difference between genotypes (Fig **3C**). Consistent with unaltered excitability, we observed no significant difference in resting membrane potential (Vrest; Fig **3D**), input resistance (Rin; Fig **3E**) or any other tested electrophysiological parameter, between genotypes (**Supplementary Table 2**).

As *in vivo* visual response reliability was significantly reduced in *Syngap* HET mice, we assessed whether L2/3 neurons displayed a lack of precision in action potential output in response to near-threshold activation, mimicking rapid synaptic activation. The amplitude of required current for near-threshold activation was 782 ± 47 pA in WT neurons, not significantly different from the 711 ± 51 pA required for *Syngap* HET mice (**Supplementary Table 2**). In response to these brief depolarising steps, we consistently observed action potential discharge (Fig **3F**) with similar latency in both WT and *Syngap* HET mice (Fig **3G**). Similarly, we found no significant difference in the jitter of action potential timing, measured as the coefficient of variation of action potential onset (CoV; Fig **3H**).

Finally, recordings of miniature excitatory postsynaptic currents (mEPSC) in the presence of tetrodotoxin (TTX) and picrotoxin (PTX) (Fig **3I**) showed no significant difference in mEPSC amplitude (Fig **3J**) and frequency (Fig **3K**) between genotypes, indicating that the strength and number of glutamatergic synapses in V1 are not affected in *Syngap* HET mice.

Taken together, these findings show that the intrinsic and synaptic properties of L2/3 pyramidal cells in adult V1 slices of *Syngap* HET mice are unaltered suggesting that differences in neuronal response reliability *in vivo* are likely due to extrinsic inputs to V1 neurons.

### Disrupted behavioural state modulation underlies decreased V1 coding precision in *Syngap* HET mice

Since intrinsic properties of V1 neurons of *Syngap* HET mice were found unaltered, we then asked whether dysregulations in behavioural state could account for the reduced reliability of visual responses in *Syngap* HET mice.

We tested whether removing behavioural state modulation through anaesthesia would improve V1 coding precision in *Syngap* HET mice. For this, we used light inhalational anaesthesia (∼0.8% isoflurane; Fig **4A**). Inhalational anaesthesia has been shown to dampen behavioural state modulation by suppressing, amongst other, both adrenergic and cholinergic inputs, while largely maintaining feedforward inputs to sensory cortical areas^46–50^. Under light anaesthesia, we found that V1 decoding accuracy for both grating orientation (Fig **4B**) and direction (Fig **4C**) was not significantly different between *Syngap* HET and WT mice (Suppl Fig **6A-B**), suggesting that feedforward inputs to V1 are unaffected in *Syngap* HET mice. On a population level, anaesthesia significantly reduced shared variance of V1 neurons for all pairs of stimulus representations (Fig **4D** **left**) and, consequently, significantly increased discriminability for both genotypes (Fig **4D** **right**). This increase was more pronounced for *Syngap* HET mice, such that V1 discriminability under anaesthesia was found to be not significantly different between genotypes (Fig **4D**).

**Figure 4.**
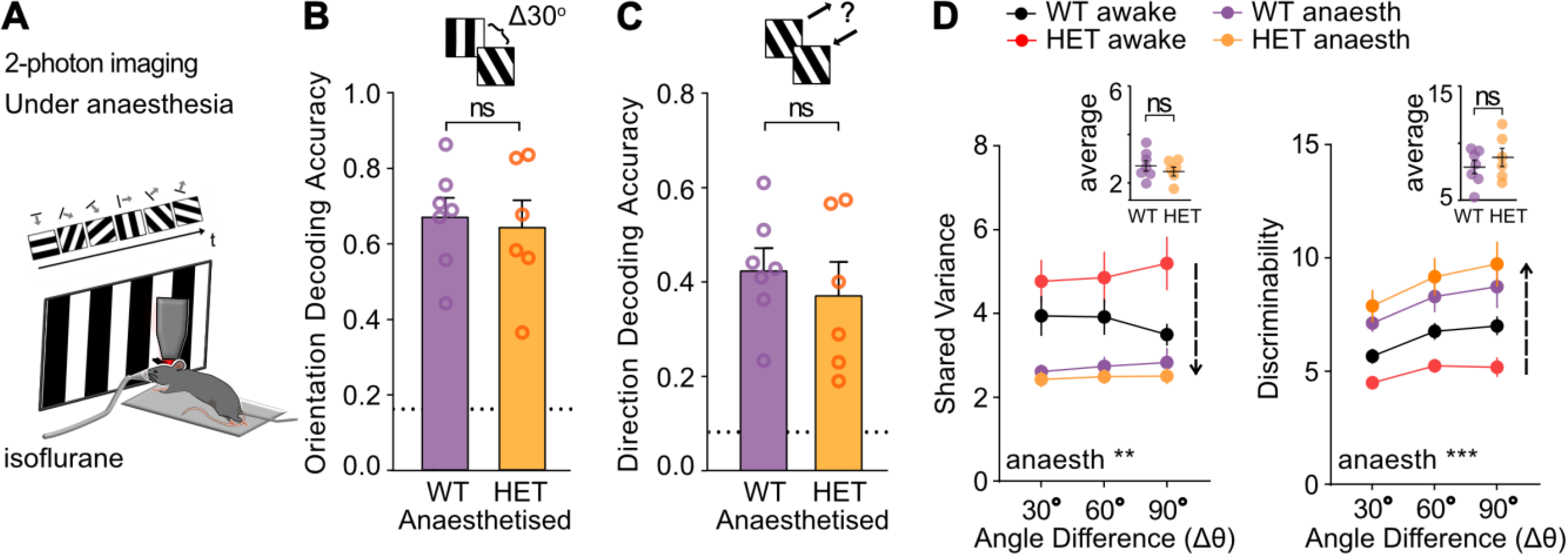
No deficits in V1 coding precision of *Syngap* HET mice under anaesthesia. **(A)** Schema of experimental method and visual stimulation under light inhalational anaesthesia (isoflurane). **(B)** Mean accuracy in decoding the correct orientation amongst 6 equally-spaced orientations (maximum likelihood decoder; *p*=0.7580, *two-tailed unpaired t-test*; n = 7 WT and n = 6 *Syngap* HET mice). Dotted line: chance level. **(C)** Mean accuracy in decoding the correct direction amongst 12 directions (maximum likelihood decoder; *p*=0.5163, *two-tailed unpaired t-test*; n = 7 WT and n = 6 *Syngap* HET mice). Dotted line: chance level. **(D) (left panel)** Anaesthesia increases discriminability of V1 neurons of *Syngap* HET mice by reducing variability of visual responses. (left panel) Mean shared variance calculated from the stimulus discriminability (anaesthesia *p*=0.0005, *REML*; n = 9 WT awake, n = 7 anaesthetised and n = 8 *Syngap* HET awake and n = 6 anaesthetised mice). Inset, mean shared variance over all angle pairs during anaesthesia (*p*=0.419, *two-tailed unpaired t-test;* n = 7 WT and n = 6 *Syngap* HET anaesthetised mice). **(D) (right panel**) Mean discriminability for pairs of gratings of set angle difference (Δθ; 30°, 60°, 90°; anaesthesia *p*<0.0001 *REML*; WT mice: n = 9 awake and n = 7 anaesthetised mice; *Syngap* HET mice: n = 8 awake and n = 6 anaesthetised mice). Inset, mean discriminability over all angle pairs during anaesthesia (*p*=0.4005, *two-tailed unpaired t-test;* n = 7 WT and n = 6 *Syngap* HET mice). The same number of neurons per animal was used to assess decoding and discriminability. Mean across animals ± SEM is shown. \**p*<0.05, \*\**p*<0.01, \*\*\**p*<0.001, \*\*\*\**p*<0.0001.

Altogether, these findings indicate that normalising behavioural state through anaesthesia increases discriminability and decoding accuracy of V1 neurons of *Syngap* HET mice by reducing variability of visual responses. These results suggest that disrupted behavioural state modulation underlies visual discrimination deficits in awake *Syngap* HET mice.

### Larger pupil size during visual stimulation in both mice and individuals with *SYNGAP1-*related disorder

Pupil diameter fluctuations can tightly track not only changes in luminance, but also variations in behavioural state such as alertness, attention, and mental engagement^51–57^. We therefore monitored pupil changes in a subset of mice (Fig **5A**). We found that pupil diameter during visual stimulation was significantly greater in *Syngap* HET mice than WT controls (Fig **5B**). This difference was not due to basal pupil resting size in darkness, which was not significantly different between genotypes (Fig **5C-D**). In addition, we found no impairment in the ability of the pupil to dilate and constrict since latency of dilation and constriction, as well as amplitude of constriction during visual stimulation, were not significantly different between *Syngap* HET and WT mice (Suppl Fig **5A-I**). These results suggest that while the pupil reflex is intact, behavioural state during visual stimulation is altered in *Syngap* HET mice compared to WT.

**Figure 5.**
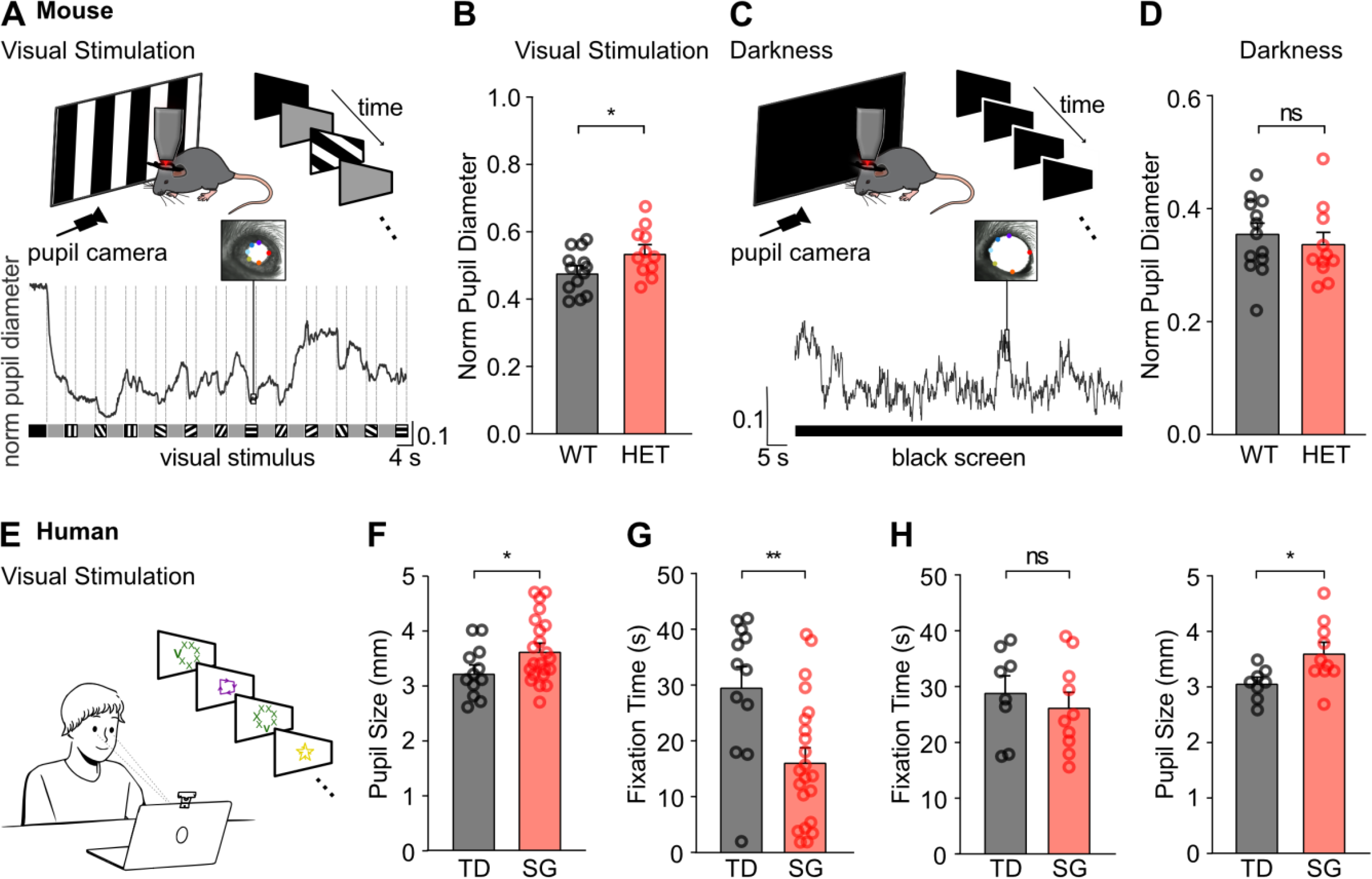
Increased pupil diameter during visual stimulation for mice and individuals with *SYNGAP1*-related disorder. **(A)** (top) Schema of experimental procedure. (bottom) Example of mouse pupil diameter trace during presentation of visual stimuli. Example image of mouse eye is shown at the point indicated on the pupil diameter trace, with coloured dots used for pupil detection through DeepLABCut. **(B)** Mean normalised mouse pupil diameter during visual stimulation (*p*=0.0479, *two-tailed unpaired t-test*; n = 13 WT and n = 11 *Syngap* HET mice). **(C)** Same as panel A during presentation of dark screen. **(D)** Mean normalised mouse pupil diameter during presentation of dark screen (*p*=0.3311, *Mann-Whitney test*; n = 13 WT and n = 11 *Syngap* HET mice). **(E)** Schema of eye tracking experiments. Visual stimuli are presented on a laptop monitor, whilst eye movements and pupil diameter are monitored using REDn Scientific eye tracking system. **(F)** Mean pupil size for typically developing (TD) and *SYNGAP1* subjects during presentation of static visual stimuli (*p*=0.0487, *two-tailed unpaired t-test*; n = 12 TD and n = 22 *SYNGAP1* individuals) **(G)** Mean total fixation time of both eyes to the screen presenting static visual stimuli (*p*=0.0029, *two-tailed unpaired t-test*; n = 12 TD and n = 22 *SYNGAP1* individuals). **(H)** (Left panel) Subsampled dataset of TD and *SYNGAP1* individuals that have comparable fixation time of both eyes to the screen (*p*=0.4962, *two-tailed unpaired t-test*; n = 8 TD and n = 10 *SYNGAP1* individuals). **(H)** (Right panel) Mean pupil size of subsampled dataset for TD and *SYNGAP1* individuals that have comparable fixation time of both eyes to the screen, as shown on the left panel (*p*=0.0249, *two-tailed unpaired t-test*; n = 8 TD and n = 10 *SYNGAP1* individuals). Mean across animals/individuals ± SEM is shown. \**p*<0.05, \*\**p*<0.01, \*\*\**p*<0.001, \*\*\*\**p*<0.0001.

To determine whether alterations in pupil size is a shared phenotype in *SYNGAP1*-related disorder, we extended our study to humans. We measured the pupil size of healthy typically developing (TD) individuals (n = 12, median age 8.3 years old) and individuals diagnosed with *SYNGAP1*-related disorder (SG; n = 22, median age 7.8 years old) while they were fixating on visual stimuli displayed on a computer screen (Fig **5E**, see also **Supplementary Table 3**). We found that SG individuals had significantly larger pupil size (Fig **5F**) than TD control individuals. One potential confounding factor was that the fixation time of some SG individuals was much lower than fixation time of TD control individuals (TD: 29.85 ± 3.5 s vs SG: 16.41 ± 2.4 s; Fig **5G**). We accounted for this difference by restricting the analysis to subjects that had comparable amounts of fixation periods (n_TD_ = 8 out of 12, n_SG_ = 10 out of 22; Fig **5H** **left**): we found that SG individuals still had larger pupil size during the presentation of visual stimuli compared to TD controls (Fig **5H** **right**).

Together, these results suggest that enlarged pupil size during visual stimulation is a shared phenotype between both mice and individuals with *SYNGAP1* haploinsufficiency, implicating altered behavioural states.

### Exogenous administration of alpha-2A noradrenergic agonist rescues V1 population coding precision in *Syngap* HET mice

Behavioural state regulation strongly depends on neuromodulatory inputs. The mouse visual cortex is densely innervated by neuromodulatory afferents, dominated by cholinergic and noradrenergic inputs^58–60^.

Noradrenaline has been shown to affect visual processing, not only by gating neuronal plasticity^61–67^ but also by regulating excitability, reliability and signal-to-noise ratios of principal neurons^54,65,68–74^ and astrocytes^75,76^ in V1. For example, local noradrenergic blockade in V1 results in hyperpolarization of the membrane potential of principal L2/3 neurons, markedly decreasing spontaneous variability^71^. We therefore tested whether modulating noradrenergic tone would rescue V1 coding precision in *Syngap* HET mice by reducing variability of visual responses. We used the α_2_-adrenergic agonist guanfacine, that dampens adrenergic tone^77^, predominately through the activation of pre-synaptic autoreceptors (α_2A_)^78^. From a translational perspective, a key advantage of using guanfacine is that it is a FDA-approved medication (FDA ref#3335794) prescribed for Attention-deficit/Hyperactivity Disorder (ADHD) and to individuals with neurodevelopmental disorders that exhibit behavioural features like sleep disturbances, anxiety-related disorders, and aggressive behaviour^79–81^.

We tested whether exogenous systemic guanfacine administration would improve V1 coding precision in *Syngap* HET mice. To avoid adverse effects like sickness and significant sedation, we used a dose of guanfacine (0.4 mg/kg, see **Methods**) previously shown to be effective at reducing hyperactivity and impulsivity in another model of neurodevelopmental disorder, the neurofibromatosis type-1 mouse model^82^. This dose of guanfacine significantly reduced open field activity in *Syngap* HET mice, resulting in similar locomotor activity (distance travelled) compared to vehicle-treated WT mice (Fig **6A, B**).

**Figure 6.**
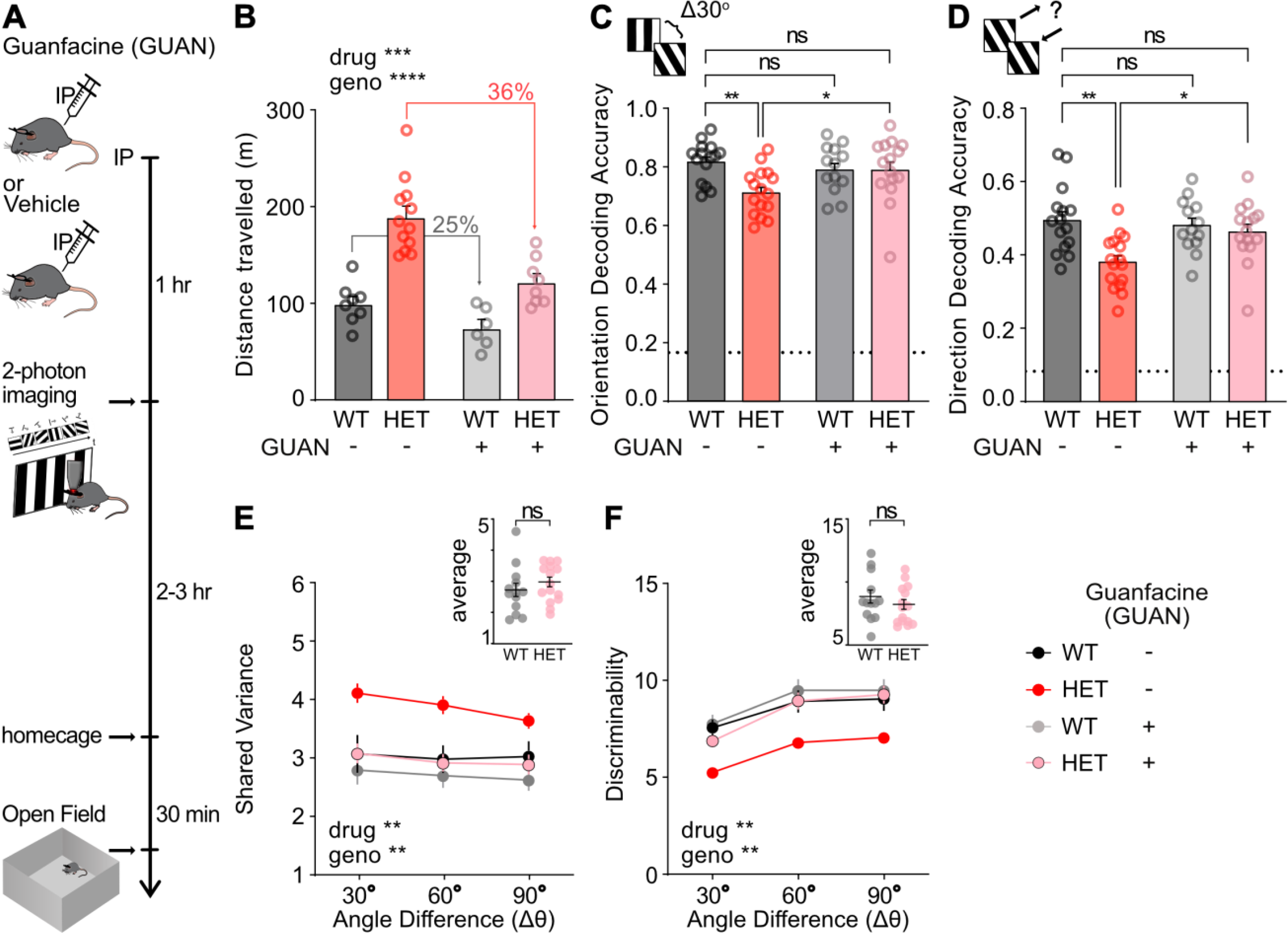
Systemic administration of guanfacine rescues V1 discriminability by reducing variance of V1 *Syngap* HET neurons. **(A)** Schema of experimental procedure. Head-fixed mice were imaged 1-1½ hrs after administration of 0.4mg/kg guanfacine (GUAN) or vehicle (saline) through intraperitoneal (IP) injection. Open field test was performed ∼30 min after the end of imaging session. **(B)** Distance travelled in open field during 30 min for control (WT) and *Syngap* HET mice after administration of vehicle (Saline; 8 WT-, 12 HET-) or 0.4 mg/kg guanfacine (GUAN; 6 WT+, 8 HET+; *2-way* ANOVA, drug *p* = 0.0001, genotype *p* < 0.0001). **(C)** Mean accuracy in decoding the correct grating orientation amongst 6 equally-spaced orientations (maximum likelihood decoder; drug x genotype *p*=0.01699, *SRH test*). Dotted line: chance level (Saline; WT-, n = 15, *Syngap* HET-, n = 16; GUAN, WT+, n =13, *Syngap* HET+, n =15 mice). **(D)** Mean accuracy in decoding the correct grating direction amongst 12 directions (maximum likelihood decoder; drug x genotype *p*=0.0466, *SRH test*). Dotted line: chance level (Saline, WT-, n = 15, *Syngap* HET-, n = 16; GUAN, WT+, n =13, *Syngap* HET+, n =15 mice). **(E)** Shared variance calculated from the stimulus discriminability (Drug *p*=0.0027, *3-way ANOVA*; Saline, WT-, n = 15, *Syngap* HET-, n = 16; GUAN, WT+, n =13, *Syngap* HET+, n =15 mice). Inset, mean variance over all angle pairs for animals that received the drug treatment (*p*=0.3374, *two-tailed unpaired t-test;* n = 13 WT and n = 15 *Syngap* HET guanfacine-treated mice). **(F)** Mean discriminability for pairs of gratings of set angle difference (Δθ; 30°, 60°, 90; Drug *p*=0.0064, *3-way ANOVA*; Saline, WT-, n = 15, *Syngap* HET-, n = 16; GUAN, WT+, n =13, *Syngap* HET+, n =15 mice). Inset, mean discriminability over all angle pairs for animals that received the drug treatment (*p*=0.3374, *two-tailed unpaired t-test;* n = 13 WT and n = 15 *Syngap* HET guanfacine-treated mice). The same number of neurons per animal was used to assess decoding and discriminability. Mean across animals ± SEM is shown. \**p*<0.05, \*\**p*<0.01, \*\*\**p*<0.001, \*\*\*\**p*<0.0001.

In V1, we found that decoding accuracy for both grating orientation (Fig **6C**) and direction (Fig **6D**) was significantly higher in guanfacine-treated *Syngap* HET mice than in vehicle-treated *Syngap* HET mice, reaching levels comparable to both guanfacine-treated and vehicle-treated WT mice (Suppl Fig **6C-D**). These results indicate that exogenous systemic guanfacine administration restored V1 coding precision deficits in *Syngap* HET mice. At a population level, guanfacine also significantly reduced the shared variance in V1 neurons across all pairs of stimuli (Fig **6E**). As a result, V1 discriminability of guanfacine-treated *Syngap* HET mice was significantly increased and not significantly different from WT levels (Fig **6F**).

These results show that targeting noradrenergic tone through systemic administration of guanfacine effectively restored V1 discriminability in *Syngap* HET mice to WT control levels.

## Discussion

Our results show that *Syngap* HET mice display significant deficits in visual discrimination, both at the level of their primary visual cortex, where population coding precision is reduced, and behaviourally, as assessed by a lower performance on a visual discrimination test. These deficits were not explained by changes in intrinsic properties of V1 neurons. In addition, V1 visual responses under anaesthesia were preserved, which indicates that visual processing impairments in awake *Syngap* HET mice are driven by a disruption of behavioural state modulation. Consistent with this hypothesis, pupil diameter during visual stimulation was found to be larger in both mice and individuals with *SYNGAP1* haploinsufficiency, implicating neuromodulatory dysfunction. Systemic administration of an α_2_-adrenoreceptor agonist restored V1 coding precision in *Syngap* HET mice to control levels. These results show that dysregulated behavioural state underlies visual discrimination deficits in *Syngap* HET mice, revealing a previously unexplored role of neuromodulatory dysregulations in *SYNGAP1*-related disorder.

### Dysregulation of neuromodulatory inputs underlies visual discrimination impairment in *Syngap* HET mice

A key finding of our study is that systemic administration of an α_2_-adrenoreceptor agonist rescued visual coding impairments, indicating a causal role of noradrenergic tone on visual disruptions in *Syngap* HET mice. This is consistent with the observation that both *Syngap* HET mice and individuals with *SYNGAP1*-related disorder exhibit larger pupil diameter during visual stimulation, since pupil diameter was shown to reflect changes in neuromodulation^56,57^. Such differences in pupil size have previously been reported in other animal models of idiopathic and monogenic forms of ASD^83^.

In *Syngap* HET mice, dysregulation of noradrenergic tone in V1 may affect visual processing through several mechanisms. *SYNGAP1* RNA expression has been reported in the human locus coeruleus (LC)^84^, the brainstem nucleus that releases noradrenaline^85–89^, suggesting that *SYNGAP1* haploinsufficiency may directly impact noradrenergic release, for example due to hyperexcitability of LC neurons. In V1, we hypothesise that increased and/or variable noradrenergic inputs lead to increased variability of V1 excitatory neuronal responses, either directly through post-synaptic activation of noradrenergic receptors, or indirectly through the modulation of inhibitory neuron activity^90–95^. Interestingly, increased gamma power in V1 has been observed in both *Syngap* HET mice and individuals with *SYNGAP1*-related disorder^96^, suggesting a potential deficit in parvalbumin-expressing (PV)-mediated inhibition^97,98^.

We propose that unreliable visual responses in V1 neurons of *Syngap* HET mice may slow the learning rate of visually-guided tasks by increasing the number of trials required to infer task rules. It has been suggested that sensory overload in ASD reduces capacity of learning^5,99^, often due to disrupting attention that causes anxiety, and physical discomfort^100^. Consistent with previous reports^101,102^, we found that *Syngap* HET mice require more trials to reach learning criterion in a visual discrimination task. This learning delay is also in agreement with other studies reporting cognitive impairments in both *Syngap* HET mice and in individuals with *SYNGAP1*-related disorder^19,103^, as well as in other neurodevelopmental disorders^35^.

### *Syngap* haploinsufficiency may have sensory modality- and cortical region-specific effects

We found intrinsic and synaptic properties of L2/3 V1 neurons to be comparable in *Syngap* HET and WT mice. This result differs from previous *in vitro* studies reporting increased excitability and synaptic function in prefrontal cortex^104,105^, hippocampus^106,107^, and primary cortical cultures^108^ but reduced excitability in primary somatosensory cortex^26^. Notably, while Michaelson et al^26^ reported reduced length and complexity of L2/3 and L4 dendrites in somatosensory cortex, we did not observe such alterations in L2/3 neurons of visual cortex. This discrepancy may explain the different mEPSC and excitability phenotypes between the two regions. *In vivo* studies also indicate differences between the somatosensory, the auditory, and the visual cortex in *Syngap* HET mice. Previous reports showed reduced responsivity of L2/3 somatosensory cortex neurons to whisker stimulation *in vivo*^26,102^. This reduced responsivity was found to persist in animals under anaesthesia^26^. By contrast, our results in visual cortex showed L2/3 V1 neurons of *Syngap* HET mice to be as responsive to visual stimuli as WT neurons. Another difference between sensory cortices was identified in a recent study using EEG to investigate sensory-evoked related potentials in *Syngap* HET mice: compared to WT controls, neural entrainment was increased in the auditory cortex but decreased in visual cortex^96^.

These results highlight modality-dependent sensory processing differences in *Syngap* HET mice, suggesting that *Syngap* haploinsufficiency has brain region specific effects. Notably, noradrenergic axons do not uniformly innervate all cortical regions^109–112^, which may further lead to cortical region-specific differences in neuromodulatory dysfunctions in *Syngap* HET mice.

### Dampening neuromodulatory tone to reduce sensory disruptions in neurodevelopmental disorders

Drugs targeting neuromodulatory systems are regularly used to address behavioural challenges in individuals with *SYNGAP1*-related disorder^80^. In the mouse model of *Syngap* haploinsufficiency, previous work has shown that the antipsychotic clozapine can partially ameliorate hyperactivity in the open field and reduce startle reactivity^113^. However, clozapine has significant side effects when used long-term^114^, including blurred vision^115^. Given these limitations, we focused on modulating noradrenergic tone in *Syngap* HET mice using systemic administration of guanfacine, an α_2_-adrenoreceptor agonist. Guanfacine is widely prescribed for Attention Deficit Hyperactivity Disorder (ADHD)^116,117^, which is increasingly co-diagnosed in individuals with *SYNGAP1*-related disorder^80,81,118^. Notably, behavioural traits observed in *Syngap* HET mice are reminiscent of ADHD symptoms in humans, including hyperactivity^103,119^, risk-taking behaviour^120^, and impulsivity during execution of sensory-guided tasks^26,121^.

Our results show that guanfacine administration restores V1 coding precision of *Syngap* HET mice to control levels, through a significant reduction in response variance of V1 HET neurons. This effect may result from guanfacine’s ability to directly suppress LC activity and, consequently, reduce norepinephrine release in the cortex. Guanfacine primarily activates pre-synaptic autoreceptors (α_2A_^78^) that dampen adrenergic tone^77,122^, by modulating both the firing rate of LC neurons and neurotransmitter release^122^. Administration of α_2_-adrenoceptor agonists to the LC has been shown to reduce firing rates in LC neurons^123–126^, to lower plasma norepinephrine concentration^127,128^, and has potent sedative effects in humans^128–131^ and rodents^132,133^. Consistent with these results, we found reduced open field locomotor activity in both *Syngap* HET and WT mice following guanfacine administration.

In addition, α_2_-adrenoceptor agonists also bind to post-synaptic receptors; albeit this effect is often considered to be masked by their presynaptic effects^122^. One cortical region that displays high expression levels of post-synaptic α_2_ receptors is the prefrontal cortex (PFC)^134^, which is also densely innervated by LC axons^109,135^. Guanfacine suppresses glutamatergic synaptic transmission and reduces excitability in the deeper layers of PFC, via predominantly post-synaptic mechanisms^136,137^. Notably, PFC of *Syngap* HET mice has been found to be more excitable than WT control, at least *in vitro*^104^. Guanfacine may therefore reduce PFC hyperexcitability and in turn affect V1 function, either through direct top-down inputs or indirectly through basalo-cortical cholinergic projection pathways^138–140^.

An important question for future research is to determine which stage of development is optimal for pharmacological intervention to ameliorate sensory dysregulations in individuals with *SYNGAP1*-related disorder^106,141–145^. In mice, *Syngap* expression in the cortex peaks early in development and gradually decreases into adulthood^146^. In the visual cortex, lower intrinsic excitability has been reported in L2/3 V1 neurons of *Syngap* HET mice during early development (P10-14)^143^, yet we find that V1 neurons in adult slices (>P60) are indistinguishable from WT control neurons. While early intervention may optimise developmental trajectories for social and cognitive impairments associated with *SYNGAP1* haploinsufficiency^143^, our results indicate that targeting noradrenergic tone through guanfacine can effectively modulate V1 circuits in adulthood. This offers the opportunity for pharmacological intervention beyond critical periods of synaptic plasticity.

Altogether, our findings reveal dysregulation of neuromodulation as a novel mechanism underlying sensory disruptions in *SYNGAP1* haploinsufficiency and highlight this as a promising therapeutic target for *SYNGAP1*-related disorder, and potentially other monogenic forms of neurodevelopmental disorders.

## Acknowledgments

We thank the GENIE Program and the Janelia Research Campus, specifically V. Jayaraman, R. Kerr, D. Kim, L. Looger, and K. Svoboda, for making GCaMP6 available. We thank N. Komiyama and S. Grant for developing and sharing the *Syngap* mouse model. We thank E. Wood for advice on the behavioural task, and R. Morris for giving us access to equipment. We thank P. Spooner for technical support. We thank P. Maeso for lab management. We thank the Bioresearch and Veterinary services staff for supporting our research by overseeing veterinary and husbandry needs. We thank all families who participated in the human study, as well as SYNGAP1 UK, the SYNGAP Research Fund and the SYNGAP1 Foundation for assisting with recruitment. This work was funded by the Simons Initiative for the Developing Brain (to D.K. and N.L.R.), the European Research Council (grant agreement 866386 to N.L.R.), the Wellcome Trust and the Royal Society (Sir Henry Dale fellowship to N.L.R.), RS MacDonald Charitable Trust (seedcorn fund to D.K.), the Shirley Foundation, the Patrick Wild Centre, the European Molecular Biology Organisation (YIP award to N.L.R.).

## Author contributions

D.K. initiated the project. D.K., N.L.R. designed the experiments. D.K. performed calcium imaging recordings. D.K., N.D. and N.K. analysed calcium imaging data. D.K. performed behavioural experiments and analysed data. D.K. performed mouse pupil recordings. D.K. and Z.C. analysed mouse pupil data. D.W. and A.K. acquired recordings of human pupil data. A.C.S. supervised the human pupil study. D.K. analysed human pupil data. S.A.B. performed *in vitro* experiments and analysed data. P.H. analysed neuronal reconstruction data. P.C.K. supervised the *in vitro* study. N.L.R. supervised the project. D.K. and N.L.R. interpreted the data, wrote, and edited the manuscript with input from all authors.

## Methods

### Animals

Animal experiments were approved by the Animal Welfare and Ethical Review Board (AWERB) of the University of Edinburgh and were performed under a project license granted by the UK Home Office and conformed with the UK Animals (Scientific Procedures) Act 1986 and the European Directive 86/609/EEC and 2010/63/EU on the protection of animals used for experimental purposes. All experiments were performed with *Syngap^+/-^*(*Syngap* HET) mutant mice that were originally generated by Komiyama *et al*. (2002)^147^, and maintained on a C57BL/6JOlaHsd ([#057]; Envigo) background line. In some experiments, *Syngap^+/-^* mice were crossed with a Cre-driver transgenic mouse line (*Pvalb*^tm1^(cre)^Arbr^/J [RRID: <u>IMSR_JAX:008069</u>] cross-bred with B6.*CgGt(ROSA)26Sor*^tm14^(CAG–tdTomato)^Hze^/J [RRID: <u>IMSR_JAX:007914</u>], Jackson Laboratory and maintained on a C57BL/6JOlaHsd [#057] Envigo background) (n=4, 1, 4 mice in Fig 1, Fig 4B-D and Suppl Fig S1-S2, respectively). All experiments were performed with *Syngap^+/+^* (*Syngap* WT) littermates as controls and with the experimenter blind to genotype. Male and female mice, aged >8 weeks, were used for experiments. Animals were typically group housed (2–5 mice) in a normal day/night 12-hour cycle.

### Participants

The human study protocol was reviewed and approved by the NHS Scotland A Research ethics committee (REC reference: 19/SS/0036). Families of participants were recruited through the Patrick Wild Centre at the University of Edinburgh with the assistance of family advocacy groups: *SYNGAP1 Foundation, SYNGAP1 UK* and *SYNGAP Research Fund* as well as through word-of-mouth. In total, 22 individuals (6 male and 16 female) who had received a diagnosis of *SYNGAP1*-related ID (SG) and 12 typically developing controls (TD; 4 male and 8 female) took part in the study. Written informed consent to participate and have their data recorded and included in the study was obtained for all participants, either from a parent, caregiver, or the participant themselves, as appropriate. Unaffected siblings that took part in the study were excluded if they had a known history of neurodevelopmental condition or an intellectual disability.

## Data acquisition

### Acute slice preparation

Acute brain slices were prepared similar to previously described^148^. Briefly, mice were anaesthetised with isoflurane, decapitated and brains rapidly removed into ice-cold, carbogenated (95 % O_2_/5 % CO_2_) sucrose-modified artificial cerebrospinal fluid (ACSF; in mM: 87 NaCl, 2.5 KCl, 25 NaHCO_3_, 1.25 NaH_2_PO_4_, 25 glucose, 75 sucrose, 7 MgCl_2_, 0.5 CaCl_2_). Once cooled, 400 μm thick slices containing the binocular region of the primary visual cortex were then cut on a Vibratome (VT1200s, Leica, Germany), then stored in sucrose-ACSF warmed to 35°C for 30 min. Slices were then transferred to room temperature until recording.

### *In vitro* whole-cell patch-clamp recordings

For electrophysiological recordings slices were transferred to a submerged recording chamber perfused with carbogenated, normal ACSF (in mM: 125 NaCl, 2.5 KCl, 25 NaHCO_3_, 1.25 NaH_2_PO_4_, 25 glucose, 1 MgCl_2_, 2 CaCl_2_) maintained at 30 ± 1°C with an inline heater (LinLab, Scientifica, UK) at a flow rate of 6-8 ml/min. Slices were visualized with IR-DIC illumination under 4x magnification (N.A. 0.1) to identify V1, then with 40x magnification to identify neurons for recording (N.A. 1.0; Olympus). Whole-cell patch-clamp recordings were made using a MultiClamp 700B amplifier (Molecular Devices, USA). Patch-pipettes were pulled from borosilicate glass capillaries (1.5 mm outer/0.86 mm inner diameter, Harvard Apparatus, UK) on a horizontal electrode puller (P-97, Sutter Instruments, CA, USA). Electrodes were filled with a K-gluconate based intracellular solution (in mM: 142 K-gluconate, 4 KCl, 0.5 EGTA, 10 HEPES, 2 MgCl_2_, 2 Na_2_ATP, 0.3 Na_2_GTP, 10 Na_2_Phosphocreatine, 2.7 Biocytin, pH=7.4, Osmolarity: 290-305 mOsm) which gave pipette resistances of 4-5 MΩ. Unless otherwise stated, all voltage-clamp recordings were performed at - 70 mV and all current-clamp recordings performed from resting membrane potential (RMP). For miniature EPSC (mEPSC) recordings, neurons were recorded as above, but with slices transferred to ACSF containing 50 µM picrotoxin (PTX) and 300 nM tetrodotoxin (TTX). Voltage-clamp recordings were performed from L2/3 principal neurons using patch-pipettes filled with an internal solution containing (in mM: 140 Cs-gluconate, 3 CsCl, 0.5 EGTA, 10 HEPES, 2 Mg-ATP, 2 Na_2_-ATP, 0.3 Na_2_-GTP, 10 Na_2_-phosphocreatine, 5 QX-314 chloride, 2.7 Biocytin), corrected to pH 7.4 with CsOH (Osm = 295 – 305 mOsm). mEPSCs were recorded for 5 min following initial wash in of the Cs-gluconate internal solution (∼2 min).

### Visualisation and reconstruction of *in vitro* recorded neurons

Following recordings, neurons were re-sealed by generating an outside-out patch configuration and then slices were immersion fixed in 4% paraformaldehyde (PFA) diluted in 0.1 M phosphate buffer (PB) at 4 °C for 24-72 hours. Visualisation was performed as previously described^149^. Slices were then transferred to phosphate buffered saline (PBS; 0.1 M phosphate buffer + 0.9% NaCl; pH: 7.4) and kept at 4 °C until processed (<3 weeks). The sections were repeatedly rinsed in PBS, then incubated in a solution containing streptavidin conjugated AlexaFluor 633 (2 µg/ml, Invitrogen, Dunfermline, UK) in PBS containing 0.5% Triton X-100 for 24-48 hours at 4 °C. Slices were then washed liberally with PBS, then PB and mounted on glass slides with hard-set mounting medium (Vectashield, Vector Labs, UK). Recorded neurons were imaged on an upright laser scanning confocal microscope (Axiovert LSM 510, Zeiss, Germany), with reconstructions performed using a 20x air-immersion objective. Z-stack images of recorded neurons were collected at 1 µm steps, with a 2048×2048 pixel image resolution.

### AAV injection and cranial window

For virus injection and cranial window implantation, 6-to 12-week-old mice were anaesthetised with isoflurane, mounted on a stereotaxic frame (David Kopf Instruments, CA), and maintained at 37 °C using a servo-driven heater (Harvard Apparatus). Non-transparent eye cream was applied to protect the eyes (Bepanthen, Bayer, Germany) and the following analgesics and anti-inflammatory drugs were injected subcutaneously pre-operatively: buprenorphine (Veteregesic; 0.1 mg kg^-1^), dexamethasone (Rapidexon; 2 μg kg^-1^) and carprofen (Carprieve; 20 mg kg^-1^). An additional injection of 25 mL kg^-1^ of warm saline Ringer’s solution (VWR, USA) was injected subcutaneously at the end of the surgery to prevent dehydration. A section of the scalp was removed, and the underlying bone was cleared from tissue and blood. A single square craniotomy (∼ 2 x 2 mm) was made over the left primary visual cortex centred at ∼3 mm lateral to midline and 1 mm anterior to lambda. After the craniotomy, adeno-associated (AAV) virus expressing the genetically encoded calcium indicator GCaMP (AAV1.Syn.GCaMP6f.WPRE.SV40; 1010-1012 IU/μL; 1:5 in aCSF; UNC, Vector Core, Chapel Hill, NC) was injected using a micromanipulator and a pipette with a 20 μm tip diameter (Nanoject, Drummond Scientific, PA) at a speed of 10nl/min, at three different depths (550, 450, and 350 μm from pia; 50-100 nl per site). Injections started at the deeper site. After each injection, the pipette was left *in situ* for an additional 5 min to prevent backflow. The craniotomy was then sealed with a custom-shaped glass coverslip (Menzel-Glaser #0) and fixed with cyano-acrylic glue. A custom-built round aluminium head-post was then implanted on the exposed skull with glue and secured in place with opaque dental acrylic (Paladur, Heraeus Kulzer, Germany). Mice were placed in a clean holding cage positioned over a heating pad and monitored until they recovered from anaesthesia, before returning to their home cage. Imaging experiments started 3-6 weeks following virus injection to allow for virus expression and clearing of the window.

### *In vivo* two-photon imaging in awake mice

Mice were placed in a cardboard tube and head-fixed in the imaging set-up. Mice were exposed to cardboard tubes in their home cage that were similar to the one present on the experimental rig, for several days prior to the imaging session. For results in Figure 1, some mice (n = 10 WT and n = 10 *Syngap* HET mice) were head-fixed in a tube while others (n = 7 WT and n = 6 *Syngap* HET mice) were head-fixed onto a cylindrical polystyrene treadmill (20 cm diameter, on a ball bearing mounting axis) and were allowed to walk freely. For results in Figures 4 and 5, all mice were head-fixed in a tube. Mice were first briefly habituated to head-fixation in the dark and during visual stimulation for 2 sessions over 2 days. Imaging was performed using a custom-built resonant scanning two-photon microscope as described previously^150^. In brief, the set up was equipped with a Ti:Sapphire excitation laser (Charmeleon Vision-S, Coherent, CA) at 920 nm and GaAsP photomultiplier tubes (Scientifica). Images were acquired at 40 Hz with a custom-programmed LabView based software (v8.2; National Instruments, UK) using an Olympus (XL PlanN 25X 1.05 NA) or a Nikon 25x water-immersion objective (Nikon; CF175 Apo 25XC W; 1.1 NA). Time-series images of one focal plane (field of view; FOV) per mouse were acquired at cortical depths between 160 and 260 μm from pia. Locomotion on the treadmill was continuously monitored using an optical encoder (250 cpr; E7P, Pewatron, Switzerland) connected to a data acquisition device (National Instruments, UK) with LabView (National Instruments, UK), sampled at 12000 Hz. Overall behaviour of the mice was monitored through an infrared camera.

### *In vivo* two-photon imaging in anaesthetised mice

In a subset of experiments after awake two-photon imaging recordings, mice were lightly sedated with chlorprothixene (1 mg/kg) and subsequently lightly anaesthetised with isoflurane (0.6 – 1 %). Eyes were covered with a thin layer of transparent eye ointment (Viscotears gel) and mice were transferred and head-fixed to the two-photon set up. Body temperature was kept constant at 37 ℃ via a closed loop temperature monitor and control system (Harvard Apparatus). Time-series images of the same focal plane as pre-anaesthesia were acquired as stated above.

### Pupil recordings during two-photon imaging in mice

Mouse pupil dilation was monitored by recording the eye contralateral to the visual stimulation screen using a camera (monochrome CMOS DMK22BUC03 or DMK22UX273 camera; ImagingSource) coupled with a fixed focal length lens (Computar M5018-MP2), positioned ∼15 cm from the mouse. Images (480 x 720) were continuously collected at 30 Hz during the imaging experiment, using the IC Capture Image Acquisition software (ImagingSource).

### Visual stimuli

Visual stimuli were generated and synchronised to the resonant scanner using the Psychophysics Toolbox package (Brainard, 1997) for MATLAB (Mathworks, MA) and displayed on an LCD monitor (51 x 29 cm, Dell, UK) placed in front of the contralateral eye (right), 20 cm from eye-level. For the habituation to head fixation in the set-up, visual stimulation consisted of full-field square-wave gratings (5 s grey screen, 5 s static, 5 s drifting, 0.05 cpd, 1 Hz, contrast 80%, mean luminance 37 cd/m^2^, 8 equally spaced directions in randomised order). On the recording day, we first performed brief retinotopic mapping to optimise the position of the screen. For recordings, visual stimuli consisted of full-field square-wave drifting gratings (2 s drifting, 0.05 cpd, 1 Hz temporal frequency, contrast 80%, mean luminance 37 cd/m^2^). Twelve equally spaced directions in randomised order were presented (30° increments), interleaved with a 4 s presentation of grey screen. 8-10 trials in total darkness and 20-25 trials during visual stimulation were presented, with dark and visual stimulation trials interleaved. For assessing pupil constriction and dilation (Suppl Fig 5), visual stimuli consisted of 5 trials of alternating black and white screen presentations (5 s each) with a 10 s of grey screen preceding that sequence. Black and white screen trials were randomly presented between visual stimulation trials with drifting gratings.

### Pharmacology

For pharmacology experiments 0.9% saline (vehicle) of guanfacine hydrochloride (GUAN; Merck) in vehicle was administered through intraperitoneal (ip) injection at a volume of 0.4 mg/kg. Drug or vehicle were delivered ∼ 1-1.5 hours prior to imaging experiments. Prior to ip injection, mice were briefly anaesthetised with isoflurane for a maximum of 20 s. Mice from the same litter with same genotype were counterbalanced for receiving either drug or vehicle. Experiments were systematically done at the same time of the light/dark cycle (first half of the light cycle) for all mice.

### Forced-choice visual discrimination task for assessing visual discrimination

Mice were trained in a two-alternative forced-choice visual discrimination task^44,45,151^. Mice were placed in a trapezoid-shaped pool (140 cm long x 80 cm wide x 40 cm high; ∼22 °C), filled with opaque water (liquid latex; Palace chemicals, Liverpool, UK). A 56 cm divider was placed perpendicular to the wider end of the pool to form 2 arms. Extra-maze cues were obscured by a white curtain. Mice had to locate a platform (10 cm diameter) that was kept invisible (∼ 2 cm submerged) in front of a specific visual cue (grating of specific orientation and direction present in one of the two arms). Visual cues were displayed on two identical computer monitors (1920 x 1080 pixels; 22-inch TK410V, LG) placed at water level at the larger side of the trapezoid pool, one associated with each arm, behind a transparent plexiglass wall (55 cm high). A black curtain was added behind the monitors. Visual cues (grey screen or square-wave gratings, 1 Hz, contrast 80%, 0.03 cpd, as seen from the edge of the platform) were generated through the psychophysics toolbox (MathWorks). Mice were tested in three phases. First (phase one), mice were trained for 4 days with a visible platform (a visible cue was placed on top of the hidden platform) (pretraining; 4 trials/day, 15 min intertrial interval (ITI)). In the second phase (visual detection; 10 trials/day, 8 days, 15 min ITI), mice were trained to swim to a hidden platform placed in the arm associated with a vertically oriented drifting grating (90°; target stimulus); the other computer screen displayed a uniform grey stimulus (non-target stimulus). The arm with the vertical grating and platform was randomized in each trial. Each trial lasted a maximum of 90 s; mice failing to find the platform were guided to the platform. If the mouse crossed the imaginary line running perpendicular to the end of the 56 cm divider (decision line) on the wrong stimulus side, the trial was recorded as an error; the mouse was left swimming until finding the platform, and then was returned immediately to the release location to perform another trial (retraining) until the mouse made a correct choice or for a maximum of 3 consecutive trials. An error trial also occurred if the mouse failed to reach any platform position within 90 s. Only mice that showed over 70% performance (average across 2 days) were used for the next phase. In the third phase (angle discrimination; 10 trials/day, 18 days, 15 min ITI), mice were trained to swim to the hidden platform placed in front of the computer screen with a vertically oriented drifting grating (90°, target stimulus) versus a 45° oriented drifting grating (non-target stimulus). All other aspects of training were the same as the second phase. In the final phase of the experiment (testing), mice were tested for their visual discrimination threshold by reducing the angle difference between the vertically oriented drifting grating (target stimulus) and the non-target stimulus. The following angle differences were tested: 30° (90° vs 60°; 10 trials/day, 2 days), 20° (90° vs 70°; 10 trials/day, 2 days), 10° (90° vs 80°; 10 trials/day, 2 days), 5° (90° vs 85°; 10 trials/day, 2 days), 25° (90° vs 65°; 10 trials/day, 2 days), 15° (90° vs 75°; 10 trials/day, 2 days). At the end of the testing phase, mice were also re-assessed in discriminating 90° vs 45° (10 trials/day, 2 days) to confirm their ability to perform the task; both groups retained >70% criterion performance (WT: 85% ± 2.1; HET: 83% ± 1.0). Finally, after the testing phase, we assessed performance for 0° angle difference (90° vs 90°, 10 trials).

All mice remained on the platform for ∼5 s before being removed from the pool. Platform locations were pseudorandomised across trials and counterbalanced across mouse groups. Release location was the same throughout all phases. Between trials, mice were dried with a towel and were returned to a holding cage (similar to their home cage) placed on a heating pad with temperature set at 37° C. A video camera mounted above the pool recorded the sessions for blind off-line analysis.

### Open field spontaneous exploration

In a subset of experiments, after two-photon imaging, mice were returned to their home cage for 30 min and were then subsequently positioned in an empty evenly lit square arena (dimensions 44 × 44 × 40 cm) made of opaque matte grey acrylic. Spontaneous exploration and locomotor activity was recorded for 30 min through a video camera mounted above the arena for blinded off-line analysis. After each mouse, the arena was extensively cleaned from droppings to eliminate odour traces.

### Eye tracking procedure and task in participants

Participants were seated on a chair approximately 60-70cm from the eye tracking laptop on which the stimuli were presented. Participants who were unable to sit alone were positioned on their caregiver’s lap, or in their own wheelchair. Stimuli were presented at 30 Hz on a 15.6-inch laptop monitor (1920 x 1080), whilst eye movements were monitored using the REDn Scientific eye tracking system (SensoMotoric Instruments GmbH; SMI), as previously described^30^. SMIs Experiment Center software was used to present stimuli and to synchronize with recorded eye movements and pupil dilation.

Prior to the start of the experiment, a five-point calibration and validation process were performed. After which, trials started with the presentation of an audio-visual animation (a moving geometric shape accompanied with a sound effect) presented for 1.5 s, which directed participants attention to the centre of the screen. Participants were then presented with an array of eight letters presented in a circle on a white background. In each array, seven of the stimuli were the letter ‘X’. The remaining letter, deemed as the target, was either a ‘+’, ‘V’, ‘S’ or an ‘O’ (odd one out). For each target, eight different arrays were created, in which the position of the target varied. Alongside this, the array letters could either be black, blue, red, or green in colour (25% for each). Each of the stimuli were presented once, in a random order, for 1.5 s. In total, there were 32 trials broken into 2 blocks. If required, short breaks were given between the blocks to allow the participants to rest and help them refocus on the task.

## Data analysis

### *Ex-vivo* whole-cell patch-clamp analysis

Voltage clamp signals were filtered at 2 kHz and current-clamp signals filtered at 5 kHz using the built in 4-pole Bessel filter of the amplifier, all digitized at 20 kHz on an analogue-digital interface (Digidata 1550B, Axon Instruments, CA, USA), and acquired with pClamp software (pClamp 10, Axon Instruments, CA, USA). Data were analysed offline using the open source Stimfit software package^152^. Cells were rejected if the RMP was > -50 mV, series resistance > 30 MΩ, or the series resistance changed by more than 20% over the course of the recording. Current-clamp recordings were performed in response to either large hyper-to depolarising current steps (-500 to +500 pA, 100 pA step, 500 ms duration, 3 repetitions) or brief depolarising steps at 25pA above rheobase current (10 ms duration, 50 repetitions).

### Neuronal reconstructions

3D reconstruction of neurons was performed using the Simple Neurite Tracer module for ImageJ/FIJI^153^, with neuronal structural properties and Sholl analysis performed from traced neurons.

### *ΔF/F_0_* acquisition

Image analysis for two-photon calcium imaging was performed as previously described^150,154^. Briefly, following image acquisition at 40Hz, brain motion was corrected with discrete Fourier 2D-based image alignment (*SIMA*^155^ 1.3.2). Regions of interest (ROIs) corresponding to cell bodies were manually segmented by inspecting down-sampled frames (2 Hz), as well as the maximum intensity projection of each imaging stack, using ImageJ/FIJI software^156^. The pixel fluorescence within each ROI was averaged to create a raw time series *F(t)*. The baseline fluorescence (*F0)* was computed by taking the fifth percentile of the smoothed *F(t)* (1-Hz low-pass, zero-phase, 60th-order FIR filter) per ROI over each trial (*F0(t)*), averaged across all trials. The *ΔF/F0* was then computed by taking the difference between *F(t)* and *F0(t)* and dividing it by *F0.* To remove neuropil contamination, we used Fast Image Signal Separation Analysis (*FISSA*^157^) which uses non-negative matrix factorization (NMF) to demix spatiotemporal overlapping signal sources through blind source separation. Python toolboxes (*SIMA* and *FISSA*) were run with WinPython 2.7.10.3 and further analysis was performed with custom-written scripts in MATLAB (MathWorks, MA).

### Stationary and Locomotion periods

Locomotion periods were defined as previously described^150,154^. Briefly, changes in the position of the cylindrical treadmill (12 KHz, interpolated onto a downsampled rate of 40Hz) were monitored and categorised based on the following criteria. Periods during which instantaneous speed was < 0.1 cm/s corresponded to stationary periods. Periods during which instantaneous speed ≥ 0.1 cm/s, 0.25 Hz low-pass filtered speed ≥ 0.1 cm/s, and an average ≥ 0.1 cm/s over a 2 s window were categorised as locomotion. Inter-locomotion intervals < 500 ms were also considered locomotion. Stationary periods < 3 s after or 0.2 s before locomotion were excluded from analysis. The locomotion modulation index (LMI) was calculated as previously described^150^ :

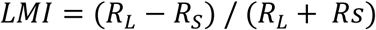

where *R*_*L*_ is the mean *ΔF/F0* during locomotion and *R*_*S*_ the mean *ΔF/F0* during stationary periods. LMI was calculated for grating-responsive neurons.

### Analysis of neuronal responses to drifting gratings

Neuronal responses to drifting gratings were defined as the mean *ΔF/F_0_* during the 2 s period of visual stimulus, subtracted by the pre-stimulus baseline, defined as the mean *ΔF/F_0_* within a 1 s window preceding the visual response. A neuron’s preferred direction (*R*_*pref*_) was defined as the drifting grating angle associated with the largest mean response.

### Grating-responsive neurons

Grating-responsive neurons were defined as those for which the mean response to their preferred direction (*R*_*pref*_) exceeded the mean response to grey screen stimuli (*R*_0_) by more than twice the standard deviation of the responses to grey stimuli (σ_0_):

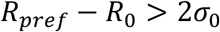

Neuronal responses to grey screen (*R*_*g*_) were defined as the mean *ΔF/F_0_* within a 2s-sliding window over the last 3s of grey screen presentation. There were no differences across genotypes in the proportion of grating-responsive neurons (*p* = 0.51, *two-tailed unpaired t-test*; n = 15 WT and n = 15 *Syngap* HET mice).

### Orientation and direction selectivity

A neuron’s selectivity to a particular drifting grating angle was quantified using the circular variance, as previously described^158^. The orientation selectivity index (OSI) and direction selectivity index (DSI) were computed as:

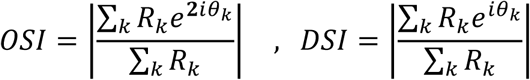

where *R*_*k*_is the average response to angle θ_*k*_, referring to either grating orientation for OSI or grating direction for DSI. OSI and DSI were calculated for grating-responsive neurons.

### Trial-to-trial reliability

The reliability of neuronal responses was quantified by measuring the correlation between all pairs of trial response vectors. A trial response vector of a given neuron was composed of its responses to the 12 grating directions. For each neuron, we calculated the correlations between all pairs of trial response vectors and applied a Fisher’s z-transformation ^159,160^. We then calculated the median of these transformed correlations, and averaged across neurons for each animal. We then reported the back-transformed average value per animal. Trial-to-trial reliability was calculated for grating-responsive neurons.

### Coefficient of Variation

The coefficient of variation was calculated as:

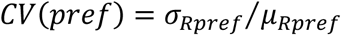

where *R*_*pref*_is the neuron’s response to the preferred direction. Coefficient of variation was calculated for grating-responsive neurons.

### Depth of modulation

The depth of modulation of each neuron was calculated as previously described^41^:

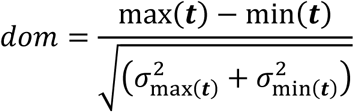

where ***t*** is the neuron’s tuning curve, obtained by averaging the responses across trials for each drifting grating stimulus, and 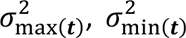 are the sample variance of the responses at the max ***t*** min ***t*** locations of the maximum (*max(t))* and minimum *(min(t)*) tuning values, respectively. Depth of modulation was calculated for grating-responsive neurons.

### Decoding of drifting grating orientation and direction

A maximum likelihood estimator (MLE) was used to decode directions based on population responses recorded with calcium imaging as previously described^161^. Briefly, decoding accuracy for each presentation of a drifting grating direction was assessed in every trial using a leave-one-out procedure. First, taking N-1 trials, the mean (μ_*k*_) and standard deviations (σ_*k*_) of the responses of every neuron associated with each presented direction (*k*) was calculated. The same number of trials (19) were used for all animals. Then, for the left-out trial, the log-likelihood of each direction *k* given the parameters μ_*k*_and σ_*k*_was calculated assuming a standard Gaussian distribution. Log-likelihoods pertaining to the same directions were summed across neurons. The direction associated with the maximum log-likelihood was selected as the decoded direction. To decode orientation, the responses of neurons to the two directions corresponding to the same orientation were first averaged. The same MLE approach was then applied to these averaged responses. Decoding accuracy was calculated as the proportion of correctly decoded orientations or directions. For each animal, we assessed performance using the same number of neurons (90 neurons; minimum number of neurons across all animals and all conditions) that were randomly sampled from the pool of all neurons 1000 times without replacement; results were averaged across samples, and finally averaged across all stimuli. Analysis was based on 20-25 trials/direction.

### Discriminability of pairs of drifting gratings

Discriminability was computed as previously described^162^. Briefly, for a given pair of two drifting grating stimuli the mean population response to each stimulus (***r***_1_ and ***r***_2_ respectively) was calculated by averaging the responses of each neuron across all trials. We determined the stimulus tuning axis Δ***f***, which is the population vector between ***r***_1_and ***r***_2_. The population responses to stimulus 1 and stimulus 2 were then projected onto the Δ***f*** axis, and for each set of projected data, mean (μ_1_ and μ_2_) and sample variance (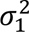 and 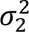) were calculated. Finally, the discriminability between the two stimuli was calculated using the d prime:

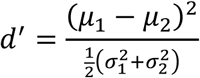

*d*^′^was computed with the same number of neurons across animals (90 neurons; minimum number of neurons across all animals and all conditions) that were randomly sampled from the pool of all neurons 1000 times without replacement; results were averaged across samples. For drifting gratings, we assessed the *d*^′^ between all pairs of directions with various increments Δ∈ {30°, ⋯, 150°}, and for each Δ angle, we averaged across all possible pairs. Before computing the *d*^′^, responses of each neuron were normalised by min-max scaling. To normalise neuronal responses (***r***), the mean response across trials (***avg***_*r*_) per grating direction was computed for each neuron. Then the minimum (*min*(***avg***_*r*_)) and maximum (*max*(***avg***_*r*_)) of these mean responses was determined. The responses were then normalised as follows:

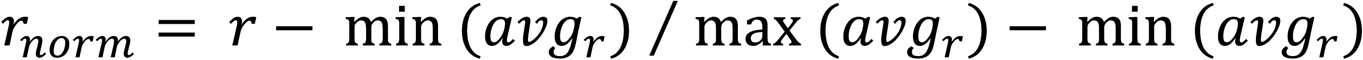

Note that *d*^′^was also computed without response normalisation and we observed the same effect (for angle pairs 30°, 60 °, 90°: genotype effect *p* = 0.0002, genotype x angle effect *p* = 0.0004; 2-way RM *ANOVA*). Analysis was based on 20-25 trials/direction.

### Behaviour

For the visual discrimination task, for training phase one (visible cue, pretraining), time to reach the visible cue from starting point was calculated averaging 4 trials each day, with no differences between genotypes (4 days; WT vs HET: 2-way RM *ANOVA*, F _(1,13)_ = 2.305, *p* = 0.1529; n = 8 WT and 7 HET mice). For training phase two (90° vs uniform grey, visual detection) performance was calculated as the percentage of correct choice (90°) across 10 trials each day, with no differences between genotypes (8 days; WT vs HET: 2-way RM *ANOVA*, F _(1,13)_ = 4.123, *p* = 0.0633; n = 8 WT and 7 HET mice). For training phase three (90° vs 45°, angle discrimination) performance was calculated as the percentage of correct choice (90°) across 10 trials each day. For the testing phase (30° to 5° angle discrimination), performance for each angle difference was calculated as the percentage of correct choices across 20 trials over 2 days (10 trials/day). For 0° angle difference, discrimination performance was calculated as the percentage of correct choices (10 trials, 1 day). Swim distance and swim speed were calculated using ANY-maze (Stoelting, Europe); calculations were based on the full path trajectory taken by the animal from the start point to the platform across all trials, irrespective of the trial outcome.

For the open field behaviour, distance travelled while exploring the open field was calculated using ANY-maze (Stoelting, Europe); calculations were based on the full path trajectory taken by the animal in the open field, during the entire time of the experiment (30 min).

### Mouse pupil data

Pupil diameter was measured from the videos of the eye using DeepLabCut^163^. For each animal 100-200 training frames were randomly sampled from the trials, making sure training frames included recordings in the dark and during visual stimulation with gratings. Six points were manually identified and spaced approximately evenly around the pupil. The network (mobileNetV2-1.0) was trained with default parameters for a maximum of 100000-200000 iterations. The pupil size was computed as the area of an ellipse fitted to the detected pupil points. If points were labelled with a likelihood of smaller than 0.9 (e.g., blinks), they were replaced with nearest-neighbour interpolation. In visual stimulation recordings, the median pupil size during visual stimulation was normalised to the 99^th^ quartile of preceding dark data. In recordings in darkness, the median pupil size was normalised to the eye size, acquired by measuring the palpebral fissure length between the endocanthion and exocanthion on ImageJ/FIJI. To calculate basic pupillary light reflex, the relative constriction was computed as the ratio of constriction amplitude and the initial pupil diameter as previously calculated^164,165^:

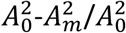

where *A_0_* is the mean pupil diameter before the onset of light reflex and *A_m_* is the minimum pupil diameter from the onset of stimulus. The constriction latency was calculated as the interval between stimulus onset and minimum pupil diameter (*A_m_*). Results were acquired by taking the mean over trials and then averaging across mice. Camera was at a fixed position during the whole duration of the experiment.

### Human pupil data

Raw measurements of average pupil diameter (in mm) were compiled in SMIs Experiment Center software as described above and exported in Microsoft Excel. Data was subsequently analysed in MATLAB. To calculate pupil diameter during visual stimulation (all types of arrays, see above), the diameter of the right eye was chosen. Both eyes had to be fixating on the screen for values to be included. Saccades, blinks, and time when only one of the two eyes were fixating were excluded from the analysis to reduce variability. Time from bouts of fixation was summed to determine total fixation time per individual. Results were acquired by taking the median over stimulation period and then averaging across participants.

### Statistics

All statistical tests, including ***n*** values and what ***n*** represents, are reported in the figure legends. Prior to group comparisons, normality (Gaussian distribution) was assessed for each data group using the D’Agostino - Pearson omnibus normality test with alpha (***α***) set at 0.05. For multiple group comparisons, two-way ANOVAs were performed. Sidak’s post-hoc tests were conducted only if the ANOVA interaction effect was significant. In case normality assumptions were violated, the data were analysed using a Scheirer-Ray-Hare (SRH) test. Since some mice imaged in awake condition were not included for the anaesthetised condition (Fig 4H and Suppl Fig S6A-B), data were analysed by fitting a mixed-effects restricted maximum likelihood model (REML). To ensure the appropriateness of the mixed-effects model, a Chi-squared test was performed prior to mixed-effects analysis of data. For single group comparisons, *t-tests* were used. In case normality assumptions were violated, Mann-Whitney U tests were used. All statistical tests were two-tailed. Unless otherwise stated, error bars in all graphs represent the standard error of the mean (SEM). Descriptive statistics and exact *p* −values can be found in **Supplementary Table 1**. In the figures, asterisks denote significant results for alpha (***α***) set at 0.05. \**p*<0.05, \*\**p*<0.01, \*\*\**p*<0.001, \*\*\*\**p*<0.0001. Statistical tests were carried out in either MATLAB or Prism 10 (GraphPad Prism).

## Figures/Figure Legends

**S1.**
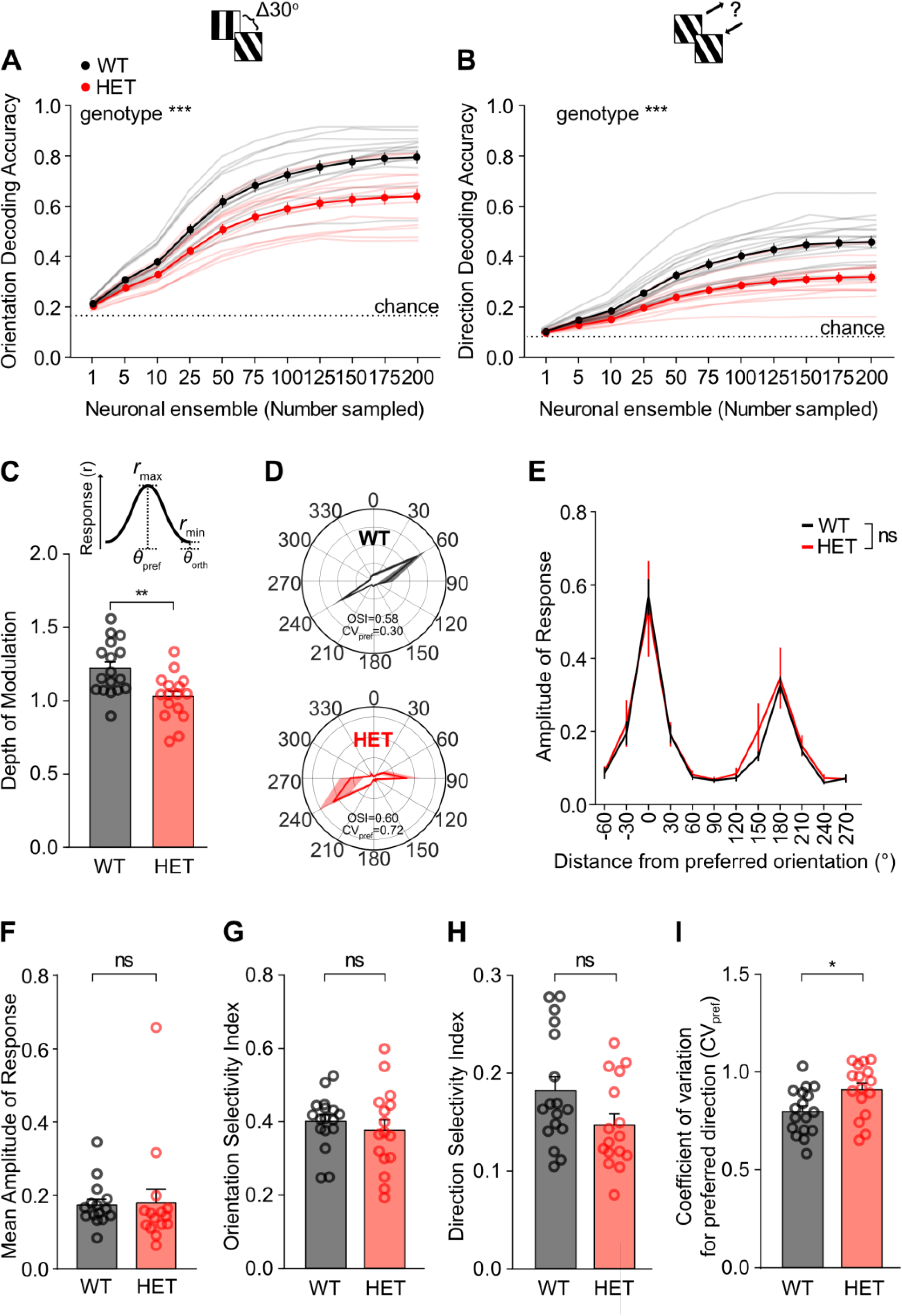
Decreased decoding accuracy for orientation and direction in V1 *Syngap* HET neurons: reduced reliability of responses but unaffected mean orientation and direction selectivity. **(A)** Decoding accuracy for grating orientation for WT and *Syngap* HET mice for randomly sampled layer 2/3 V1 neuronal ensembles (1-200 neurons; WT vs HET *p*=0.0009, *RM 2-way ANOVA*; n = 15 WT and n = 15 *Syngap* HET mice). Dotted line: chance level. Light coloured lines, individual animals. Bold lines, mean and SEM across animals. **(B)** Same as panel A for grating direction (1-200 neurons; WT vs HET *p*=0.0003, *RM 2-way ANOVA*; n = 15 WT and n = 15 *Syngap* HET mice). **(C)** Mean depth of modulation (*p*=0.0038, *two-tailed unpaired t-test*; n = 15 WT and n = 15 *Syngap* HET mice). **(D)** Example polar plots of visual responses from one WT and one *Syngap* HET V1 neuron. Polar plots show *ΔF/F_0_* values, the radius is 0.2 *ΔF/F_0_*. Bold line represents average responses over trials. Shaded error bar represents SEM over trials. OSI: Orientation Selectivity Index, CV_pref_: Coefficient of variation of response to preferred direction. **(E)** Mean direction tuning curves, aligned to the response to the preferred direction (WT vs HET *p*=0.6987, *RM 2-way ANOVA*; n = 15 WT and n = 15 *Syngap* HET mice). **(F)** Mean amplitude of response averaged across all directions (*p*=0.2017, *Mann-Whitney test*; n = 15 WT and n = 15 *Syngap* HET mice). **(G)** Mean Orientation Selectivity Index (OSI; *p*=0.38, *two-tailed unpaired t-test*; n = 15 WT and n = 15 *Syngap* HET mice). **(H)** Mean Direction Selectivity Index (DSI; *p*=0.0601, *two-tailed unpaired t-test*; n = 15 WT and n = 15 *Syngap* HET mice). **(I)** Mean coefficient of variation of response to the preferred direction (CV_pref_; *p*=0.0219, *two-tailed unpaired t-test*; n = 15 WT and n = 15 *Syngap* HET mice). Mean across animals ± SEM is shown. \**p*<0.05, \*\**p*<0.01, \*\*\**p*<0.001, \*\*\*\**p*<0.0001.

**S2.**
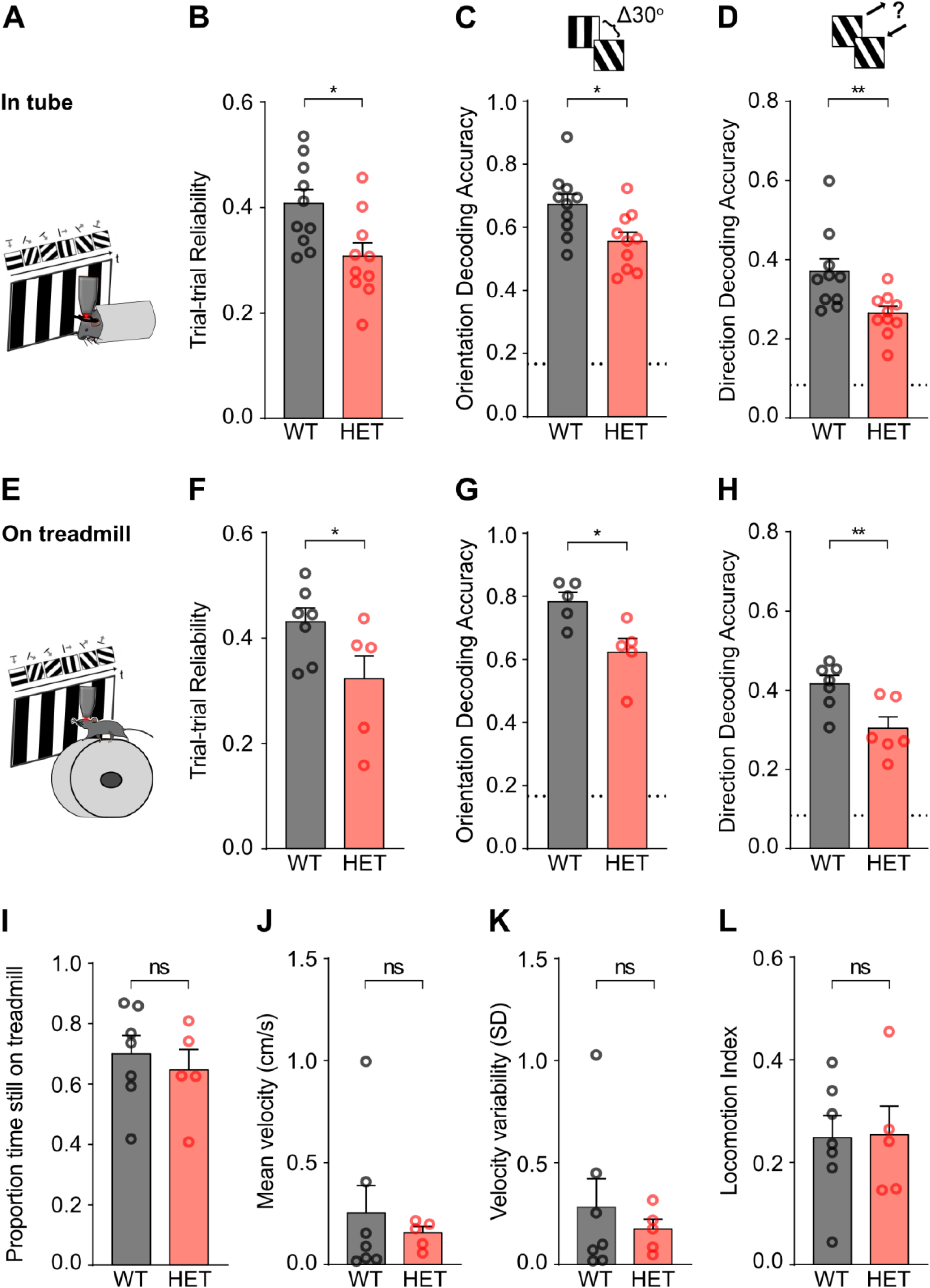
Decreased reliability of visual responses and coding precision of V1 neurons of *Syngap* HET mice are not due to differences in locomotion. **(A)** Schema of experimental procedure for head-fixed mice placed in a tube. **(B)** Mean trial-to-trial reliability of visual responses for WT and *Syngap* HET mice (*p*=0.0129, *two-tailed unpaired t-test*; n = 10 WT and n = 10 *Syngap* HET mice). **(C)** Mean accuracy in decoding grating orientation amongst 6 equally-spaced orientations (maximum likelihood decoder; *p*=0.0151, *two-tailed unpaired t-test*; n = 10 WT and n = 10 *Syngap* HET mice). Dotted line: chance level. **(D)** Same as panel C for grating direction (maximum likelihood decoder; *p*=0.0039, *Mann-Whitney test*; n = 10 WT and n = 10 *Syngap* HET mice). Dotted line: chance level. **(E)** Schema of experimental procedure for head-fixed mice placed on a cylindrical treadmill. **(F)** Same as B for mice on treadmill (*p*=0.0484, *two-tailed unpaired t-test*; n = 5 WT and n = 5 *Syngap* HET mice). **(G)** Same as C for mice on treadmill (maximum likelihood decoder; *p*=0.017, *two-tailed unpaired t-test*; n = 5 WT and n = 5 *Syngap* HET mice). **(H)** Same as D for mice on treadmill (maximum likelihood decoder; *p*=0.0137, *two-tailed unpaired t-test*; n = 5 WT and n = 5 *Syngap* HET mice). **(I)** Proportion of time spent still on the treadmill during the presentation of visual stimuli (*p*=0.3051, *two-tailed unpaired t-test*; n = 5 WT and n = 4 *Syngap* HET mice). Note that locomotion speed was not collected for 1 *Syngap* HET mouse (deficient optical encoder) included in panels F-H. **(J)** Mean self-initiated locomotion velocity during the presentation of visual stimuli (in cm/s; *p*=0.5913, *two-tailed unpaired t-test*; n = 5 WT and n = 4 *Syngap* HET mice). **(K)** Standard deviation of self-initiated locomotion velocity over trials (*p*=0.4839, *two-tailed unpaired t-test*; n = 5 WT and n = 4 *Syngap* HET mice). **(L)** Mean Locomotion modulation index during visual stimulation (LMI; *p*=0.7254, *two-tailed unpaired t-test*; n = 5 WT and n = 4 *Syngap* HET mice). Mean across animals ± SEM is shown. \**p*<0.05, \*\**p*<0.01, \*\*\**p*<0.001, \*\*\*\**p*<0.0001.

**S3.**
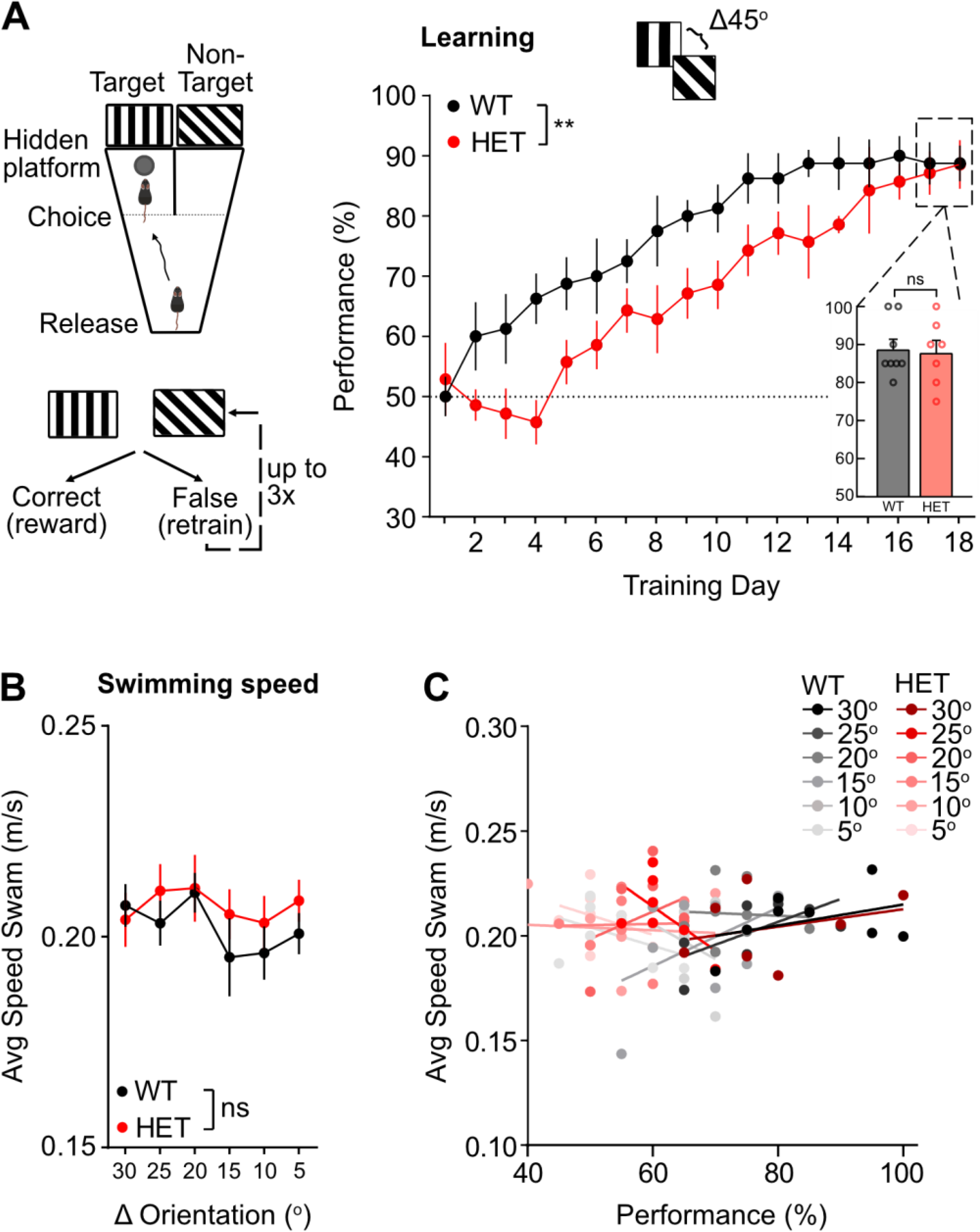
Learning of visual discrimination task and swimming speed. **(A)** Schema of experimental procedure. (top) Mice were placed in a modified water Y-maze and had to choose the arm associated with a vertical drifting target grating (90°) to reach a hidden platform and escape the water. (bottom) Learning paradigm. If the target arm is chosen, trial is classified as correct, and animal is returned to holding cage. If non-target arm is chosen, trial is classified as incorrect, and animal is positioned back at the release location to repeat the trial (retraining), for up to 3 consecutive trials. (Right panel), Learning of coarse discrimination (45°). Behavioural performance of WT and *Syngap* HET mice (WT vs HET *p*=0.0081, *RM 2-way ANOVA*; n = 8 WT and n = 7 *Syngap* HET mice) on day 1 to day 18 of learning. Inset: Mean performance of last two days of learning (*p*=0.8324, *two-tailed unpaired t-test*; n = 8 WT and n = 7 *Syngap* HET mice). Inset same as Figure 2B. **(B)** Mean speed swam during the testing of fine discrimination shown in Figure 2C (WT vs HET *p*=0.525, *RM 2-way ANOVA*; n = 8 WT and n = 7 *Syngap* HET mice). **(C)** Mean speed swam during all trials as a function of behavioural performance. Each point is the mean speed swam by a single animal averaged across trials for a given discrimination difficulty (angle difference between target and non-target gratings of 30°, 25°, 20°, 15°, 10°, and 5°) plotted against performance at that discrimination difficulty (n= 8 WT and 7 *Syngap* HET). Bold lines: regression lines. See **Supplementary Table 1** for extended statistics. Mean across animals ± SEM is shown. \**p*<0.05, \*\**p*<0.01, \*\*\**p*<0.001, \*\*\*\**p*<0.0001.

**S4.**
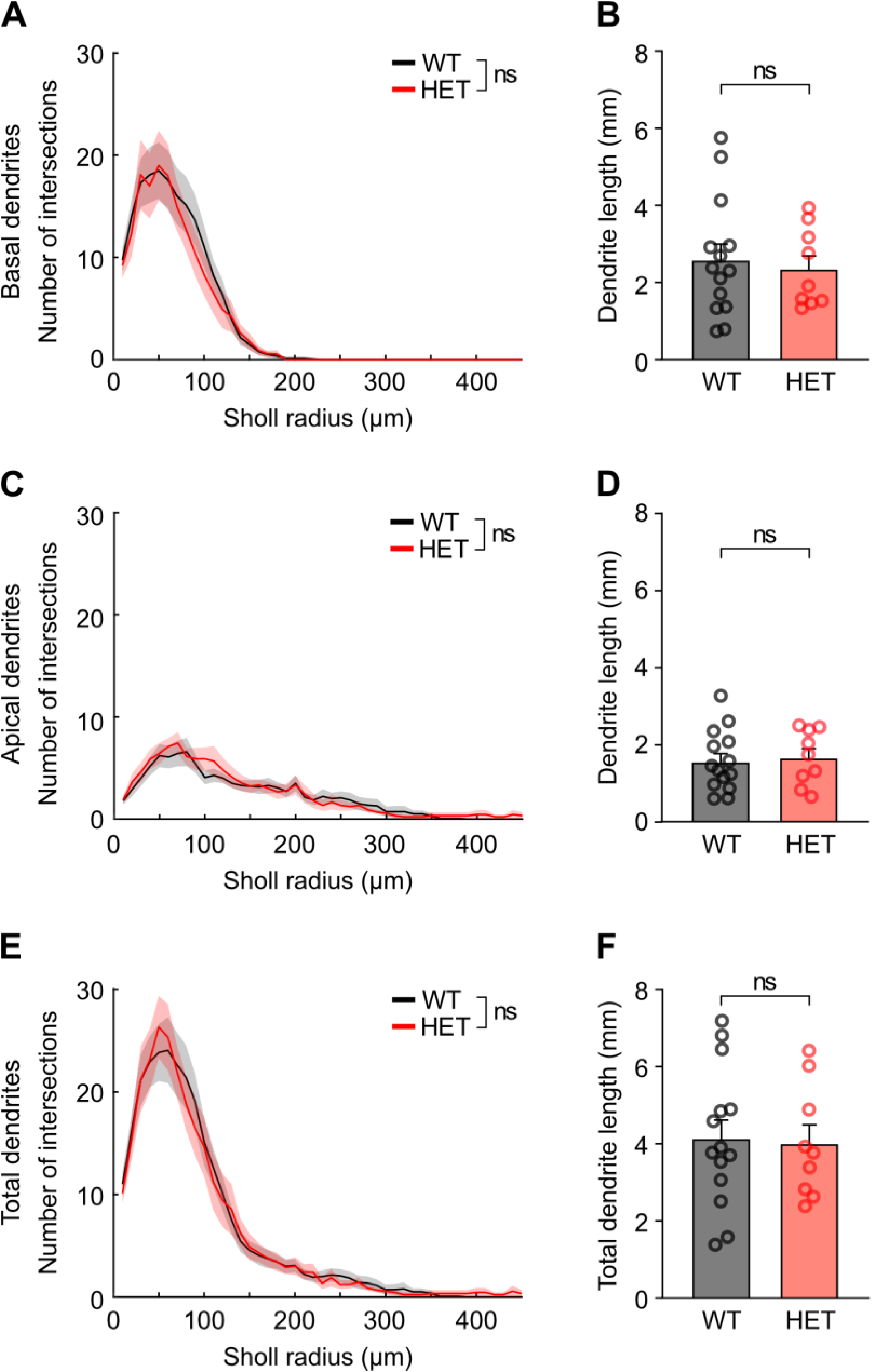
L2/3 pyramidal neurons in V1 are morphologically similar in both *Syngap* HET and WT mice. **(A)** Sholl distribution of basal dendrites from reconstructed L2/3 pyramidal cells from WT and *Syngap* HET mice (WT vs HET *p*=0.2806, *RM 2-way ANOVA*; n = 15 WT and n = 9 *Syngap* HET mice). **(B)** Quantification of the length of basal dendrites from reconstructed L2/3 pyramidal cells from WT and *Syngap* HET mice (*p*=0.691, *two-tailed unpaired t-test*). **(C)** Sholl distribution of apical dendrites from reconstructed L2/3 pyramidal cells from WT and *Syngap* HET mice (WT vs HET *p*=0.2882, *RM 2-way ANOVA*; n = 15 WT and n = 9 *Syngap* HET mice). **(D)** Quantification of the length of apical dendrites from reconstructed L2/3 pyramidal cells from WT and *Syngap* HET mice (*p*=0.758, *two-tailed unpaired t-test*). **(E)** Sholl distribution of total dendritic domain from reconstructed L2/3 pyramidal cells from WT and *Syngap* HET mice (WT vs HET *p*=0.7338, *RM 2-way ANOVA*; n = 15 WT and n = 9 *Syngap* HET mice). **(F)** Quantification of total dendritic length from reconstructed L2/3 pyramidal cells from WT and *Syngap* HET mice (*p*=0.855, *two-tailed unpaired t-test*). Mean across animals ± SEM is shown. \**p*<0.05, \*\**p*<0.01, \*\*\**p*<0.001, \*\*\*\**p*<0.0001.

**S5.**
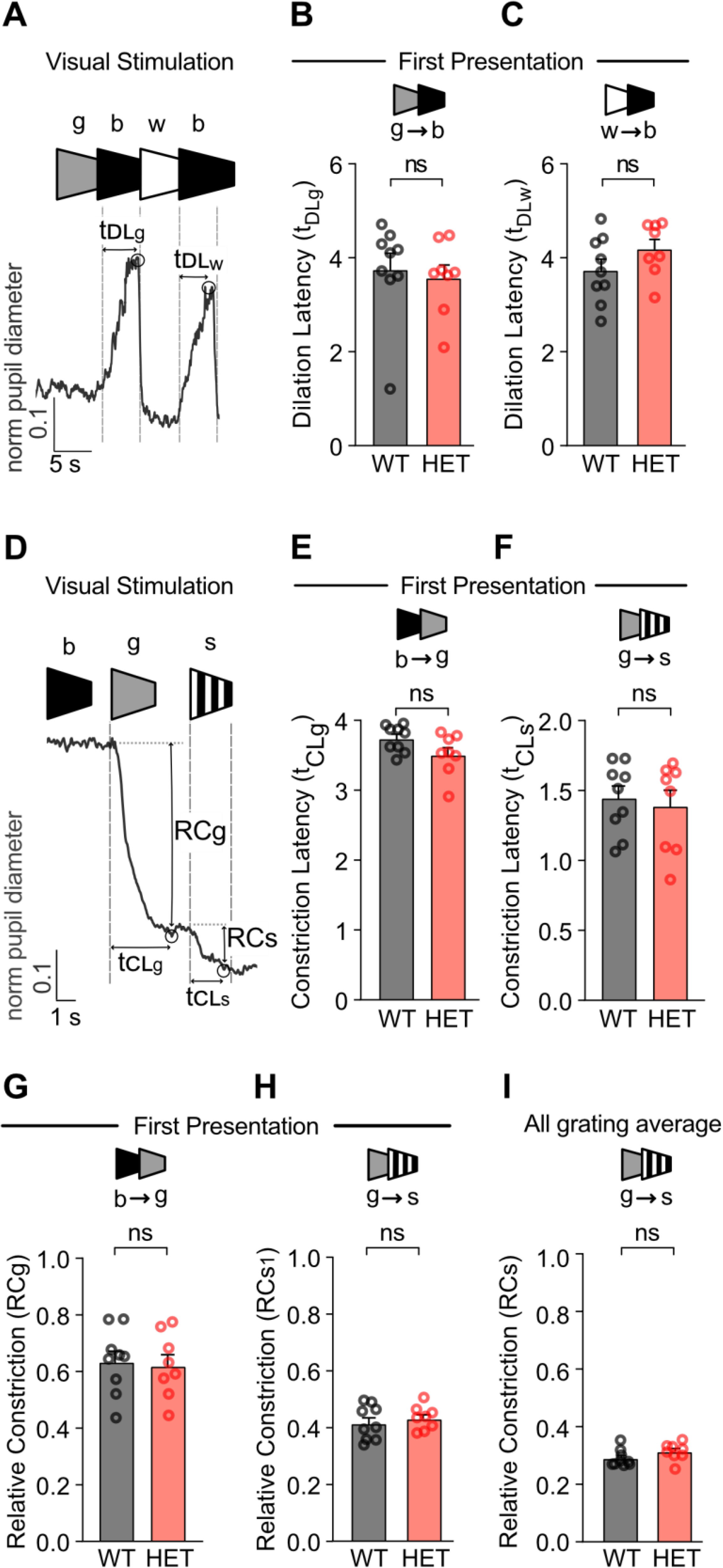
Intact pupil reflex in *Syngap* HET mice: no impairment in the capacity of the pupil to dilate and constrict. **(A)** Example trace of pupil diameter during visual stimulation and illustration of pupillary light reflex parameters: dilation latency from grey (t_DLg_) and dilation latency from white (t_DLw_) (see ***Methods***). Presentation of grey stimulus for 10 s, black 5 s, and white 5 s. **(B)** Mean dilation latency of pupil diameter for the first transition from grey to black stimulus (t_DLg_; *p*=0.5593, *Mann-Whitney test*; n = 9 WT and n = 8 *Syngap* HET mice). **(C)** Mean dilation latency of pupil diameter for the first transition from white to black stimulus (t_DLw_; *p*=0.1670, *two-tailed unpaired t-test*; n = 9 WT and n = 8 *Syngap* HET mice). **(D)** Example trace of pupil diameter and illustration of pupillary light reflex parameters: constriction latency during grey (t_CLg_), constriction latency during grating stimulus (t_CLs_:), relative constriction during grey (RC_g_), relative constriction during grating stimulus (RC_s_) (see ***Methods***). Presentation of grey stimulus for 4 s and grating stimulus 2 s. **(E)** Mean constriction latency of pupil diameter during the first transition from black to grey stimulus (t_CLg_; *p*=0.0702, *two-tailed unpaired t-test*; n = 9 WT and n = 8 *Syngap* HET mice). **(F)** Mean constriction latency of pupil diameter during the first transition from grey to grating stimulus (t_CLs_; *p*=0.6798, *two-tailed unpaired t-test*; n = 9 WT and n = 8 *Syngap* HET mice). **(G)** Mean relative constriction of pupil diameter during the first transition from black to grey stimulus (RC_g_; *p*=0.8010, *two-tailed unpaired t-test*; n = 9 WT and n = 8 *Syngap* HET mice). **(H)** Mean relative constriction of pupil diameter during the first transition from grey to grating (RC_s1_; *p*=0.5292, *two-tailed unpaired t-test*; n = 9 WT and n = 8 *Syngap* HET mice). **(I)** Mean relative constriction of pupil diameter for transitions from grey to grating (RC_s_; *p*=0.1289, *two-tailed unpaired t-test*; n = 9 WT and n = 8 *Syngap* HET mice). Mean across animals ± SEM is shown. \**p*<0.05, \*\**p*<0.01, \*\*\**p*<0.001, \*\*\*\**p*<0.0001.

**S6.**
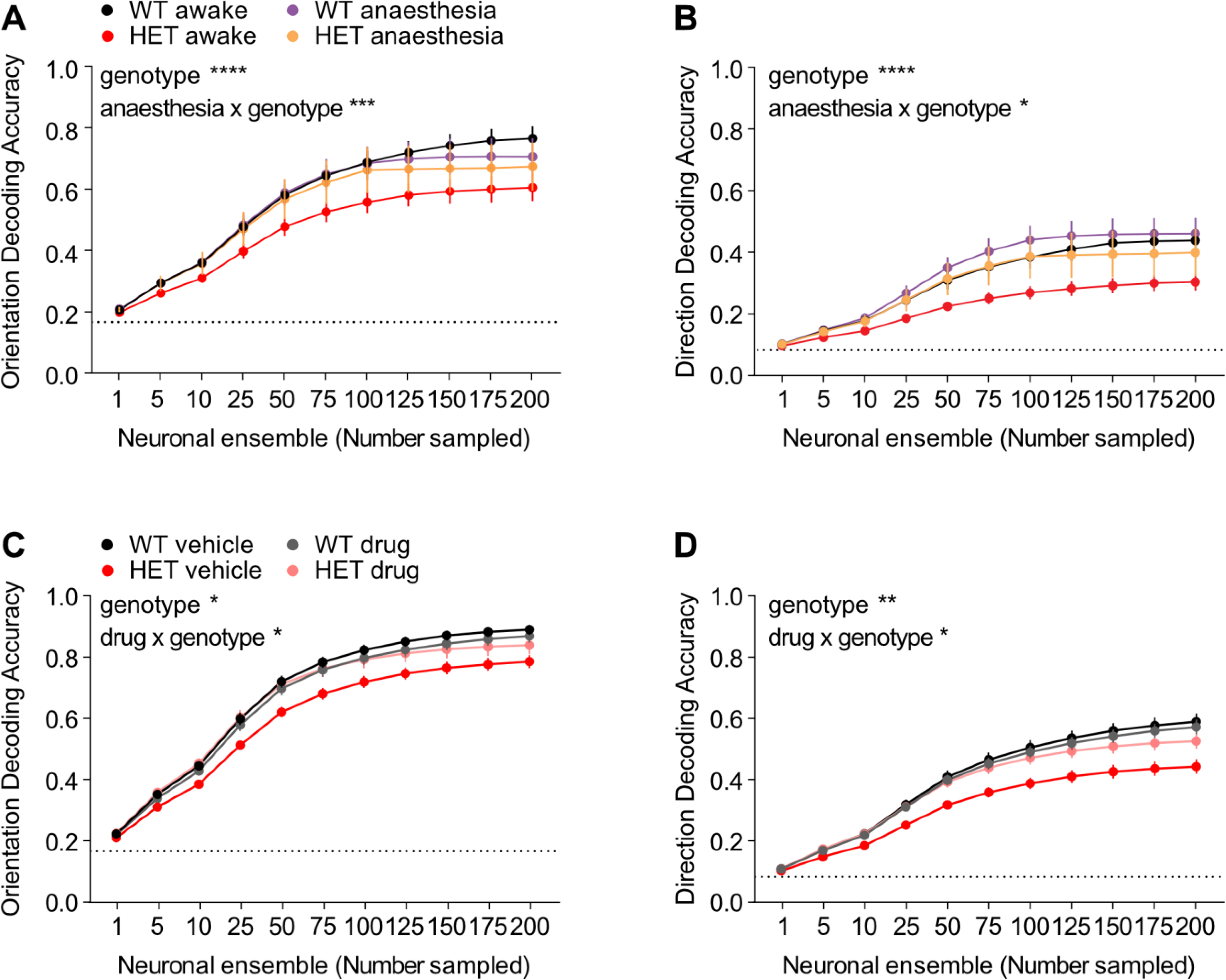
Anaesthesia and guanfacine rescue V1 coding precision deficits in *Syngap* HET mice. **(A)** Decoding accuracy for grating orientation for randomly sampled layer 2/3 V1 neuronal ensembles (1-200 neurons) in awake and anesthetized WT and *Syngap* HET mice (genotype *p*<0.0001, genotype x anaesthesia *p*=0.0005, *REML*; n = 9 WT awake, n = 7 anaesthetised and n = 8 *Syngap* HET awake and n = 6 anaesthetised mice). Dotted line: chance level. **(B)** Same as panel A for grating direction (genotype *p*<0.0001, genotype x anaesthesia *p*=0.0269, *REML*; n = 9 WT awake, n = 7 anaesthetised and n = 8 *Syngap* HET awake and n = 6 anaesthetised mice). **(C)** Decoding accuracy for grating orientation for randomly sampled layer 2/3 V1 neuronal ensembles (1-200 neurons) in WT and *Syngap* HET mice injected with either saline (vehicle) or 0.4mg/kg guanfacine (drug) (genotype *p*=0.0227, genotype x drug *p*=0.0224, *3-way ANOVA*; saline: WT – n = 15, HET – n = 16, GUAN: WT + n =13, HET + n =15 mice). Dotted line: chance level. **(D)** Same as panel C for grating direction (genotype *p*=0.0019, genotype x drug *p*=0.0263, *3-way ANOVA*; saline: WT – n = 15, HET – n = 16, GUAN: WT + n =13, HET + n =15 mice). Mean across animals ± SEM is shown. \**p*<0.05, \*\**p*<0.01, \*\*\**p*<0.001, \*\*\*\**p*<0.0001.

## Supplementary Tables

**Table S1.**
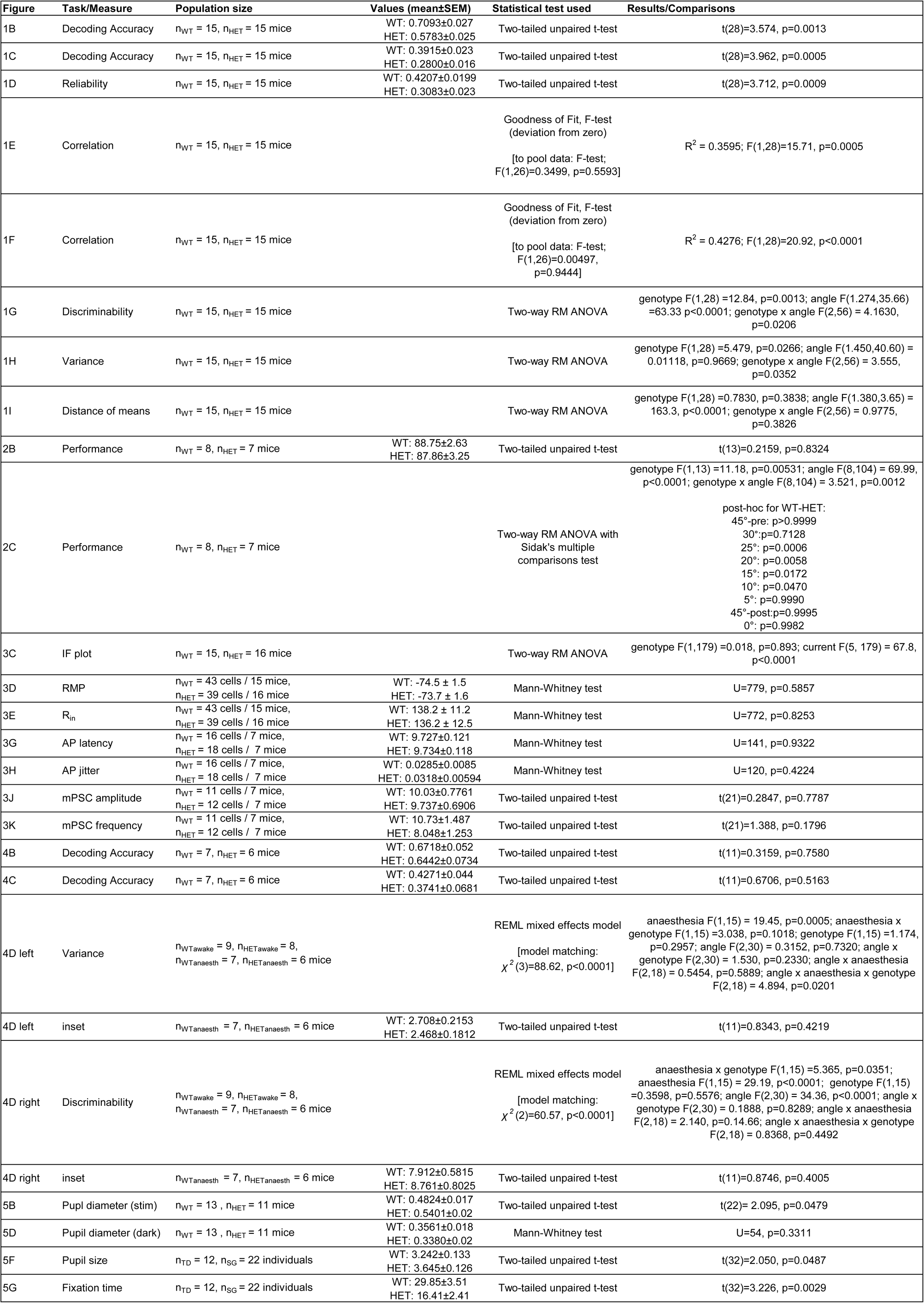
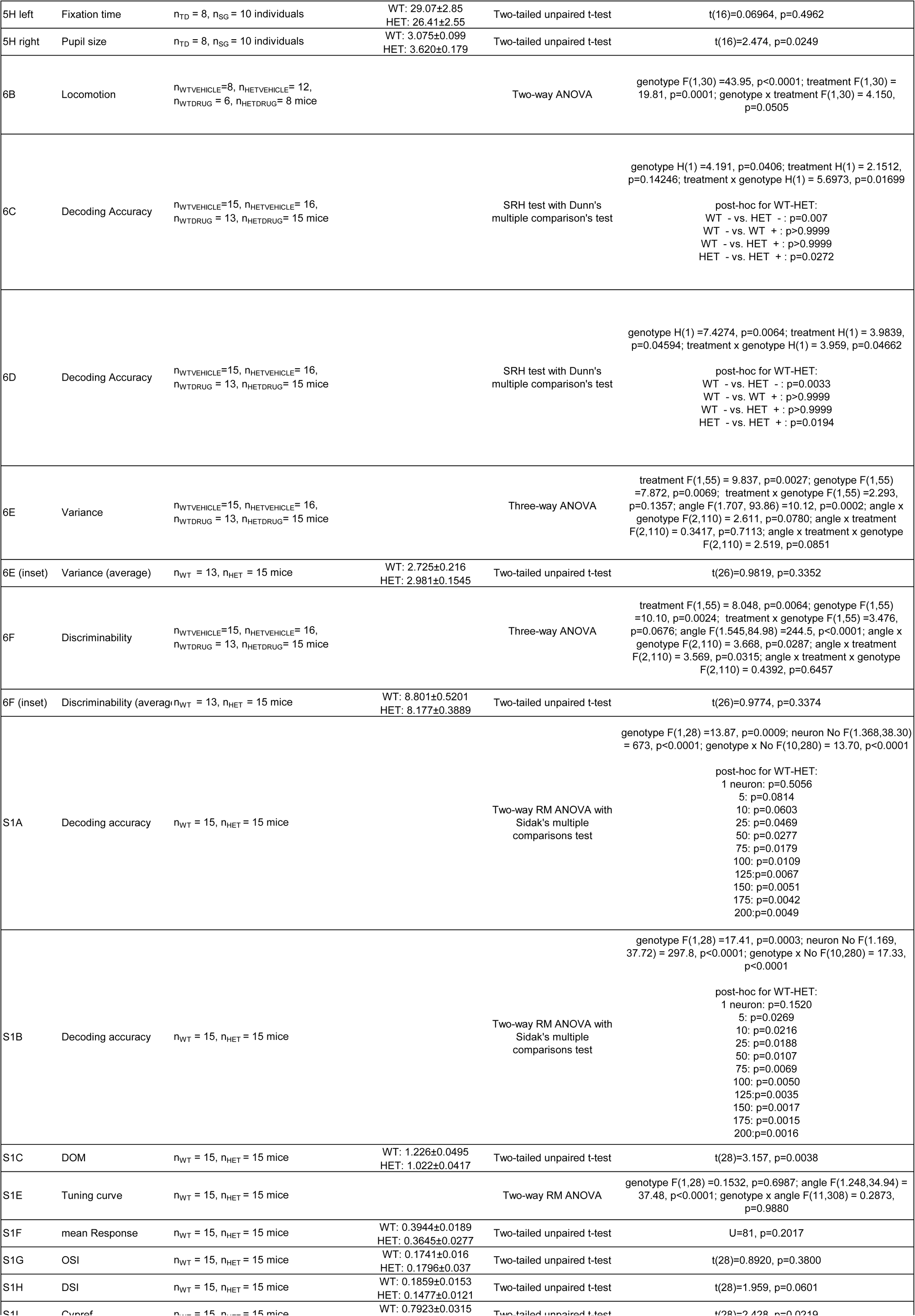
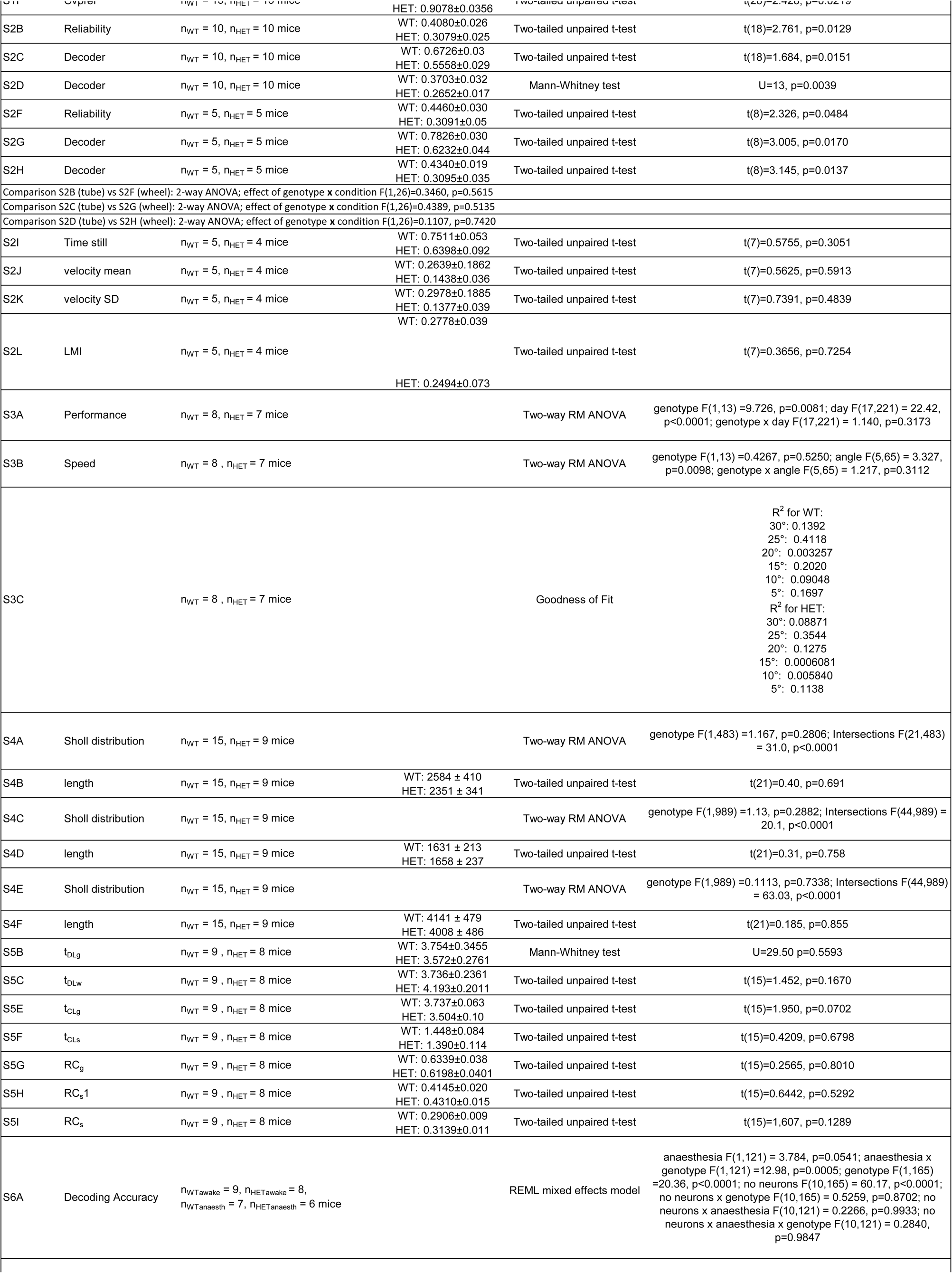
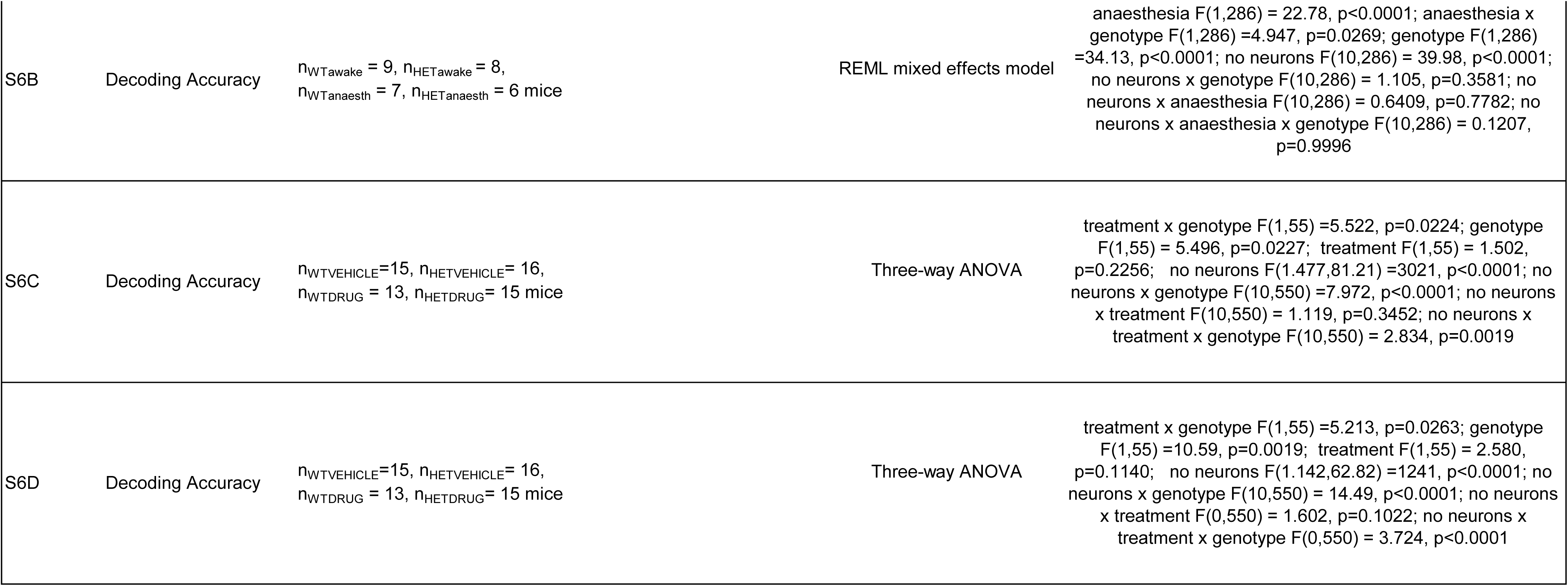
Statistical tests and results by figure panel. Separate excel document.

**Table 2:**
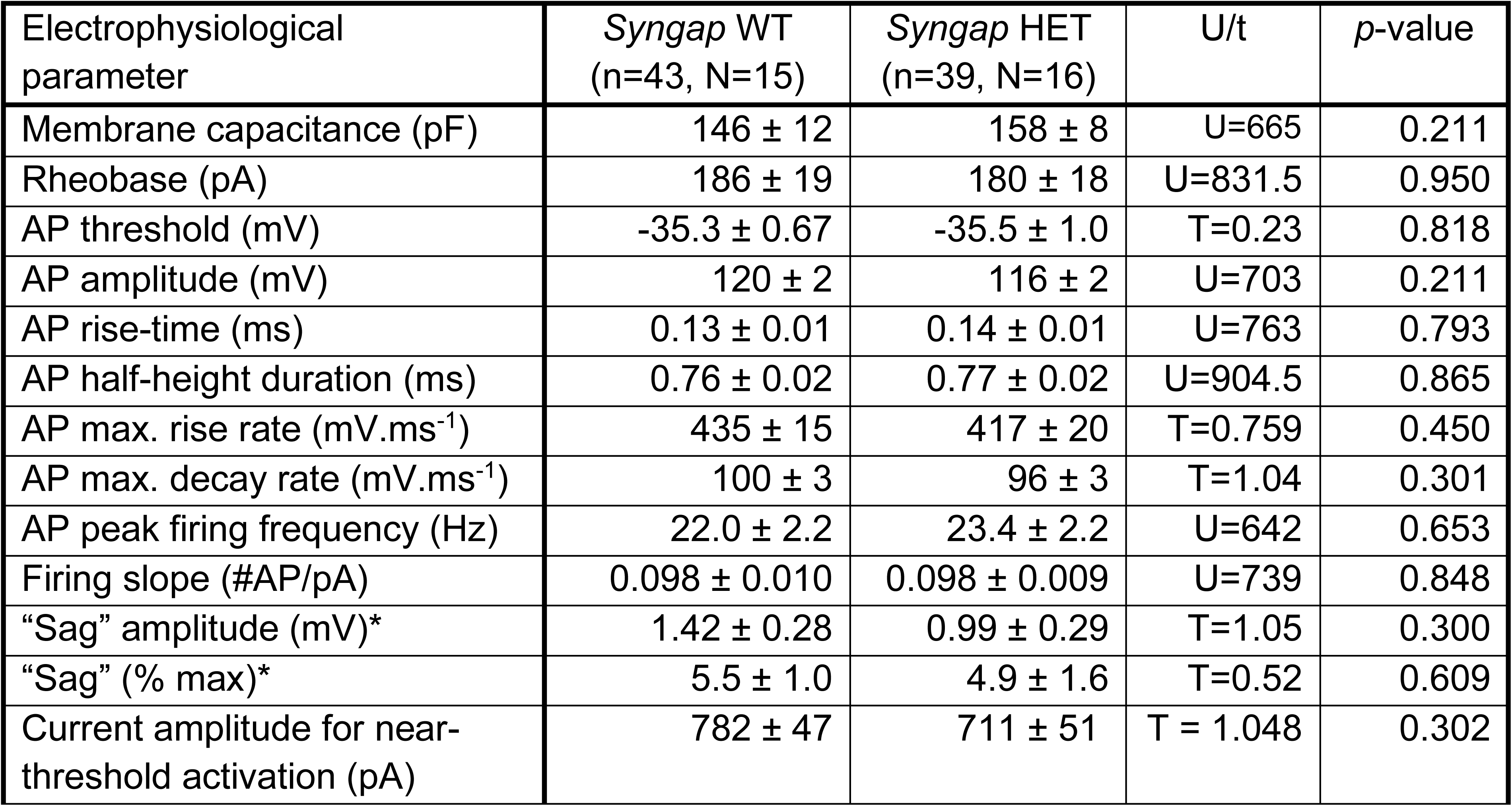
Summary of L2/3 pyramidal neuron physiological parameters in *Syngap* HET and WT mice. Data for intrinsic physiological parameters are shown as mean ± SEM. n represents number of neurons, N represents number of animals. * Note for “sag” values the number of measurements differ: WT, n=24 neurons in N=8 mice and *Syngap* HET, n=17 neurons in N=8 mice.

**Table 3.**
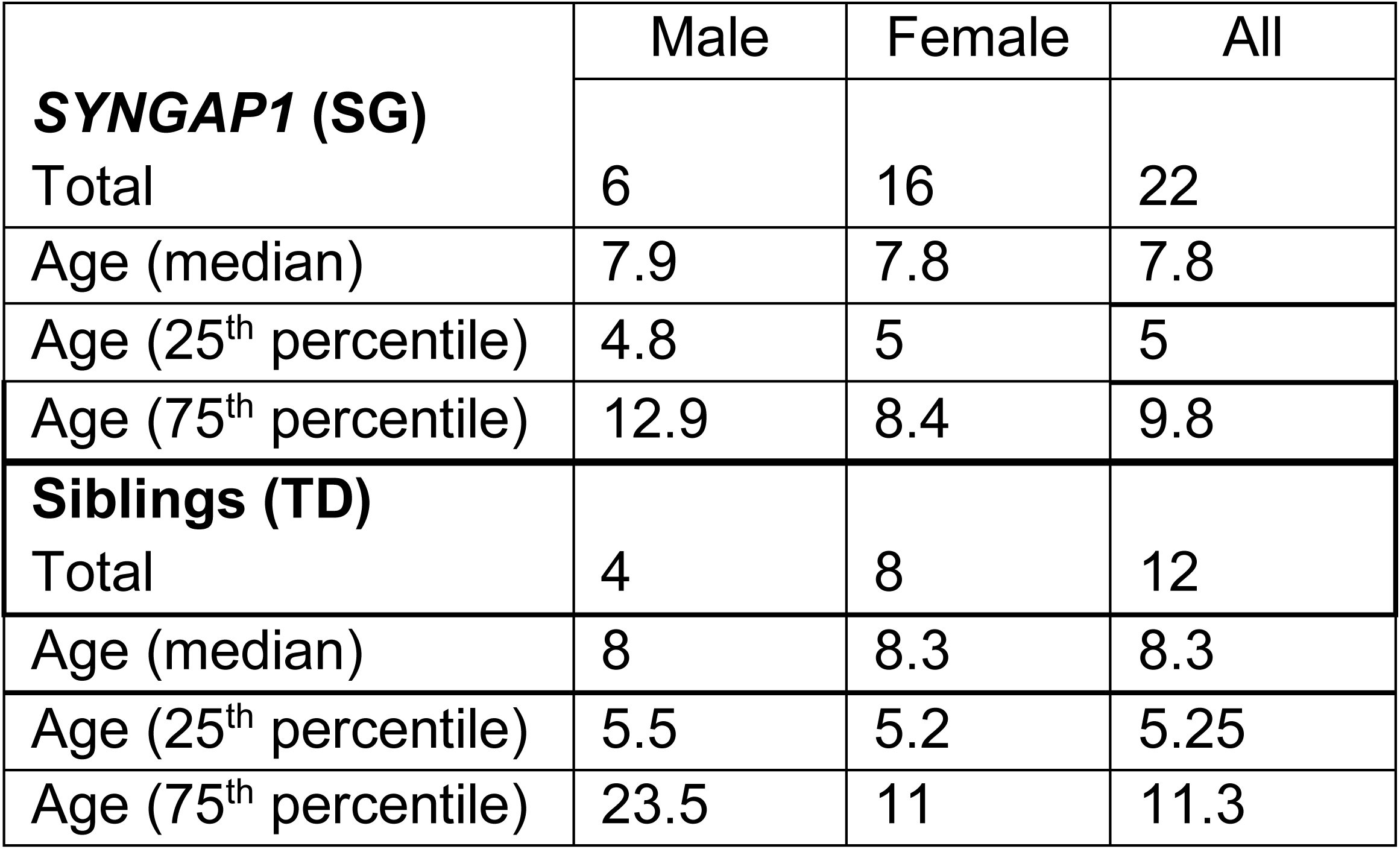
Description of participants included in. **Figure 5**. Table lists the demographics of the *SYNGAP1* (SG) and typically developing (TD) sibling populations recruited for this study. Age in years.

## References

1. Diagnostic and Statistical Manual of Mental Disorders DSM Library. https://dsm.psychiatryonline.org/doi/book/10.1176/appi.books.9780890425596.

2. Estes, A., Zwaigenbaum, L., Gu, H., St John, T., Paterson, S., Elison, J.T., Hazlett, H., Botteron, K., Dager, S.R., Schultz, R.T., et al. (2015). Behavioral, cognitive, and adaptive development in infants with autism spectrum disorder in the first 2 years of life. J Neurodev Disord 7, 24. 10.1186/s11689-015-9117-6.

3. Robertson, C.E., and Baron-Cohen, S. (2017). Sensory perception in autism. Nat Rev Neurosci 18, 671–684. 10.1038/nrn.2017.112.

4. Marco, E.J., Hinkley, L.B.N., Hill, S.S., and Nagarajan, S.S. (2011). Sensory Processing in Autism: A Review of Neurophysiologic Findings. Pediatr Res 69, 48–54. 10.1203/PDR.0b013e3182130c54.

5. Joosten, A.V., and Bundy, A.C. (2010). Sensory processing and stereotypical and repetitive behaviour in children with autism and intellectual disability. Australian Occupational Therapy Journal 57, 366–372. 10.1111/j.1440-1630.2009.00835.x.

6. Satterstrom, F.K., Kosmicki, J.A., Wang, J., Breen, M.S., De Rubeis, S., An, J.-Y., Peng, M., Collins, R., Grove, J., Klei, L., et al. (2020). Large-Scale Exome Sequencing Study Implicates Both Developmental and Functional Changes in the Neurobiology of Autism. Cell 180, 568–584.e23. 10.1016/j.cell.2019.12.036.

7. Deciphering Developmental Disorders Study (2015). Large-scale discovery of novel genetic causes of developmental disorders. Nature 519, 223–228. 10.1038/nature14135.

8. McRae, J.F., Clayton, S., Fitzgerald, T.W., Kaplanis, J., Prigmore, E., Rajan, D., Sifrim, A., Aitken, S., Akawi, N., Alvi, M., et al. (2017). Prevalence and architecture of de novo mutations in developmental disorders. Nature 542, 433–438. 10.1038/nature21062.

9. Hamdan, F.F., Gauthier, J., Spiegelman, D., Noreau, A., Yang, Y., Pellerin, S., Dobrzeniecka, S., Côté, M., Perreau-Linck, E., Carmant, L., et al. (2009). Mutations in SYNGAP1 in autosomal nonsyndromic mental retardation. N Engl J Med 360, 599–605. 10.1056/NEJMoa0805392.

10. Chen, H.-J., Rojas-Soto, M., Oguni, A., and Kennedy, M.B. (1998). A Synaptic Ras-GTPase Activating Protein (p135 SynGAP) Inhibited by CaM Kinase II. Neuron 20, 895–904. 10.1016/S0896-6273(00)80471-7.

11. Kim, J.H., Lee, H.-K., Takamiya, K., and Huganir, R.L. (2003). The Role of Synaptic GTPase-Activating Protein in Neuronal Development and Synaptic Plasticity. J. Neurosci. 23, 1119–1124. 10.1523/JNEUROSCI.23-04-01119.2003.

12. Kim, J.H., Liao, D., Lau, L.-F., and Huganir, R.L. (1998). SynGAP: a Synaptic RasGAP that Associates with the PSD-95/SAP90 Protein Family. Neuron 20, 683–691. 10.1016/S0896-6273(00)81008-9.

13. A role for synGAP in regulating neuronal apoptosis - Knuesel - 2005 - European Journal of Neuroscience - Wiley Online Library https://onlinelibrary.wiley.com/doi/full/10.1111/j.1460-9568.2005.03908.x.

14. Mastro, T.L., Preza, A., Basu, S., Chattarji, S., Till, S.M., Kind, P.C., and Kennedy, M.B. (2020). A sex difference in the response of the rodent postsynaptic density to synGAP haploinsufficiency. eLife 9, e52656. 10.7554/eLife.52656.

15. Katsanevaki, D., Till, S.M., Buller-Peralta, I., Nawaz, M.S., Louros, S.R., Kapgal, V., Tiwari, S., Walsh, D., Anstey, N.J., Petrović, N.G., et al. (2024). Key roles of C2/GAP domains in SYNGAP1-related pathophysiology. Cell Reports 43. 10.1016/j.celrep.2024.114733.

16. Parker, M.J., Fryer, A.E., Shears, D.J., Lachlan, K.L., McKee, S.A., Magee, A.C., Mohammed, S., Vasudevan, P.C., Park, S., Benoit, V., et al. (2015). De novo, heterozygous, loss-of-function mutations in SYNGAP1 cause a syndromic form of intellectual disability. Am J Med Genet A 167, 2231–2237. 10.1002/ajmg.a.37189.

17. J Lloyd Holder, J., Hamdan, F.F., and Michaud, J.L. (2019). SYNGAP1-Related Intellectual Disability. In GeneReviews® [Internet] (University of Washington, Seattle).

18. Agarwal, M., Johnston, M.V., and Stafstrom, C.E. (2019). *SYNGAP1* mutations: Clinical, genetic, and pathophysiological features. International Journal of Developmental Neuroscience 78, 65–76. 10.1016/j.ijdevneu.2019.08.003.

19. Gamache, T.R., Araki, Y., and Huganir, R.L. (2020). Twenty Years of SynGAP Research: From Synapses to Cognition. J. Neurosci. 40, 1596–1605. 10.1523/JNEUROSCI.0420-19.2020.

20. Hamdan, F.F., Gauthier, J., Araki, Y., Lin, D.-T., Yoshizawa, Y., Higashi, K., Park, A.-R., Spiegelman, D., Dobrzeniecka, S., Piton, A., et al. (2011). Excess of de novo deleterious mutations in genes associated with glutamatergic systems in nonsyndromic intellectual disability. Am J Hum Genet 88, 306–316. 10.1016/j.ajhg.2011.02.001.

21. Berryer, M.H., Hamdan, F.F., Klitten, L.L., Møller, R.S., Carmant, L., Schwartzentruber, J., Patry, L., Dobrzeniecka, S., Rochefort, D., Neugnot-Cerioli, M., et al. (2013). Mutations in SYNGAP1 cause intellectual disability, autism, and a specific form of epilepsy by inducing haploinsufficiency. Hum Mutat 34, 385–394. 10.1002/humu.22248.

22. Mignot, C., von Stülpnagel, C., Nava, C., Ville, D., Sanlaville, D., Lesca, G., Rastetter, A., Gachet, B., Marie, Y., Korenke, G.C., et al. (2016). Genetic and neurodevelopmental spectrum of SYNGAP1-associated intellectual disability and epilepsy. J Med Genet 53, 511–522. 10.1136/jmedgenet-2015-103451.

23. Wright, D., Kenny, A., Mizen, L.A.M., McKechanie, A.G., and Stanfield, A.C. (2024). The Behavioral Profile of SYNGAP1-Related Intellectual Disability. American Journal on Intellectual and Developmental Disabilities 129, 199–214. 10.1352/1944-7558-129.3.199.

24. Weldon, M., Kilinc, M., Lloyd Holder, J., and Rumbaugh, G. (2018). The first international conference on SYNGAP1-related brain disorders: a stakeholder meeting of families, researchers, clinicians, and regulators. Journal of Neurodevelopmental Disorders 10, 6. 10.1186/s11689-018-9225-1.

25. Wright, D., Kenny, A., Eley, S., McKechanie, A.G., and Stanfield, A.C. (2022). Clinical and behavioural features of SYNGAP1-related intellectual disability: a parent and caregiver description. Journal of Neurodevelopmental Disorders 14, 34. 10.1186/s11689-022-09437-x.

26. Michaelson, S.D., Ozkan, E.D., Aceti, M., Maity, S., Llamosas, N., Weldon, M., Mizrachi, E., Vaissiere, T., Gaffield, M.A., Christie, J.M., et al. (2018). SYNGAP1 Heterozygosity Disrupts Sensory Processing by Reducing Touch-Related Activity within Somatosensory Cortex Circuits. Nat Neurosci 21, 1–13. 10.1038/s41593-018-0268-0.

27. Prchalova, D., Havlovicova, M., Sterbova, K., Stranecky, V., Hancarova, M., and Sedlacek, Z. (2017). Analysis of 31-year-old patient with SYNGAP1 gene defect points to importance of variants in broader splice regions and reveals developmental trajectory of SYNGAP1-associated phenotype: case report. BMC Med Genet 18, 62. 10.1186/s12881-017-0425-4.

28. von Stülpnagel, C., Hartlieb, T., Borggräfe, I., Coppola, A., Gennaro, E., Eschermann, K., Kiwull, L., Kluger, F., Krois, I., Møller, R.S., et al. (2019). Chewing induced reflex seizures (“eating epilepsy”) and eye closure sensitivity as a common feature in pediatric patients with SYNGAP1 mutations: Review of literature and report of 8 cases. Seizure 65, 131–137. 10.1016/j.seizure.2018.12.020.

29. Vlaskamp, D.R.M., Shaw, B.J., Burgess, R., Mei, D., Montomoli, M., Xie, H., Myers, C.T., Bennett, M.F., XiangWei, W., Williams, D., et al. (2019). SYNGAP1 encephalopathy: A distinctive generalized developmental and epileptic encephalopathy. Neurology 92, e96–e107. 10.1212/WNL.0000000000006729.

30. Wright, D., Kenny, A., Eley, S., McKechanie, A.G., and Stanfield, A.C. (2024). Visual social attention in SYNGAP1-related intellectual disability. Autism Research 17, 1083–1093. 10.1002/aur.3148.

31. Dunn, W. Sensory Profile 2.User’s Manual; Pearson: New York, NY, USA, 2014.

32. Lyons-Warren, A.M., McCormack, M.C., and Holder, J.L. (2022). Sensory Processing Phenotypes in Phelan-McDermid Syndrome and SYNGAP1-Related Intellectual Disability. Brain Sci 12, 137. 10.3390/brainsci12020137.

33. Damianidou, E., Mouratidou, L., and Kyrousi, C. (2022). Research models of neurodevelopmental disorders: The right model in the right place. Front. Neurosci. 16. 10.3389/fnins.2022.1031075.

34. Bhaskaran, A.A., Gauvrit, T., Vyas, Y., Bony, G., Ginger, M., and Frick, A. (2023). Endogenous noise of neocortical neurons correlates with atypical sensory response variability in the Fmr1-/y mouse model of autism. Nat Commun 14, 7905. 10.1038/s41467-023-43777-z.

35. Goel, A., Cantu, D.A., Guilfoyle, J., Chaudhari, G.R., Newadkar, A., Todisco, B., de Alba, D., Kourdougli, N., Schmitt, L.M., Pedapati, E., et al. (2018). Impaired perceptual learning in a mouse model of Fragile X syndrome is mediated by parvalbumin neuron dysfunction and is reversible. Nat Neurosci 21, 1404–1411. 10.1038/s41593-018-0231-0.

36. Chen, Q., Deister, C.A., Gao, X., Guo, B., Lynn-Jones, T., Chen, N., Wells, M.F., Liu, R., Goard, M.J., Dimidschstein, J., et al. (2020). Dysfunction of cortical GABAergic neurons leads to sensory hyper-reactivity in a Shank3 mouse model of ASD. Nat Neurosci 23, 520–532. 10.1038/s41593-020-0598-6.

37. Townsend, L.B., Jones, K.A., Dorsett, C.R., Philpot, B.D., and Smith, S.L. (2020). Deficits in higher visual area representations in a mouse model of Angelman syndrome. J Neurodev Disord 12, 28. 10.1186/s11689-020-09329-y.

38. Auerbach, B.D., Manohar, S., Radziwon, K., and Salvi, R. (2021). Auditory hypersensitivity and processing deficits in a rat model of fragile X syndrome. Neurobiol Dis 161, 105541. 10.1016/j.nbd.2021.105541.

39. Wallace, M.L., van Woerden, G.M., Elgersma, Y., Smith, S.L., and Philpot, B.D. (2017). Ube3a loss increases excitability and blunts orientation tuning in the visual cortex of Angelman syndrome model mice. J Neurophysiol 118, 634–646. 10.1152/jn.00618.2016.

40. Montijn, J.S., Vinck, M., and Pennartz, C.M.A. (2014). Population coding in mouse visual cortex: response reliability and dissociability of stimulus tuning and noise correlation. Frontiers in Computational Neuroscience 8.

41. Chettih, S.N., and Harvey, C.D. (2019). Single-neuron perturbations reveal feature-specific competition in V1. Nature 567, 334–340. 10.1038/s41586-019-0997-6.

42. Dadarlat, M.C., and Stryker, M.P. (2017). Locomotion Enhances Neural Encoding of Visual Stimuli in Mouse V1. J. Neurosci. 37, 3764–3775. 10.1523/JNEUROSCI.2728-16.2017.

43. Averbeck, B.B., and Lee, D. (2006). Effects of Noise Correlations on Information Encoding and Decoding. Journal of Neurophysiology 95, 3633–3644. 10.1152/jn.00919.2005.

44. Padamsey, Z., Katsanevaki, D., Dupuy, N., and Rochefort, N.L. (2022). Neocortex saves energy by reducing coding precision during food scarcity. Neuron 110, 280–296.e10. 10.1016/j.neuron.2021.10.024.

45. Prusky, G.T., West, P.W.R., and Douglas, R.M. (2000). Behavioral assessment of visual acuity in mice and rats. Vision Research 40, 2201–2209. 10.1016/S0042-6989(00)00081-X.

46. Constantinople, C.M., and Bruno, R.M. (2011). Effects and Mechanisms of Wakefulness on Local Cortical Networks. Neuron 69, 1061–1068. 10.1016/j.neuron.2011.02.040.

47. Kushikata, T., Yoshida, H., Kudo, M., Kudo, T., Kudo, T., and Hirota, K. (2011). Role of coerulean noradrenergic neurones in general anaesthesia in rats. British Journal of Anaesthesia 107, 924–929. 10.1093/bja/aer303.

48. Violet, J.M., Downie, D.L., Nakisa, R.C., Lieb, W.R., and Franks, N.P. (1997). Differential Sensitivities of Mammalian Neuronal and Muscle Nicotinic Acetylcholine Receptors to General Anesthetics. Anesthesiology 86, 866–874. 10.1097/00000542-199704000-00017.

49. Flood, P., Ramirez-Latorre, J., and Role, L. (1997). Alpha4beta2 Neuronal Nicotinic Acetylcholine Receptors in the Central Nervous System Are Inhibited by Isoflurane and Propofol, but alpha7-type Nicotinic Acetylcholine Receptors Are Unaffected. Anesthesiology 86, 859–865. 10.1097/00000542-199704000-00016.

50. Raz, A., Grady, S.M., Krause, B.M., Uhlrich, D.J., Manning, K.A., and Banks, M.I. (2014). Preferential effect of isoflurane on top-down vs. bottom-up pathways in sensory cortex. Frontiers in Systems Neuroscience 8.

51. Iriki, A., Tanaka, M., and Iwamura, Y. (1996). Attention-induced neuronal activity in the monkey somatosensory cortex revealed by pupillometrics. Neuroscience Research 25, 173–181. 10.1016/0168-0102(96)01043-7.

52. Hoeks, B., and Levelt, W.J.M. (1993). Pupillary dilation as a measure of attention: a quantitative system analysis. Behavior Research Methods, Instruments, & Computers 25, 16–26. 10.3758/BF03204445.

53. Kahneman, D., and Beatty, J. (1966). Pupil diameter and load on memory. Science 154, 1583– 1585. 10.1126/science.154.3756.1583.

54. McGinley, M.J., Vinck, M., Reimer, J., Batista-Brito, R., Zagha, E., Cadwell, C.R., Tolias, A.S., Cardin, J.A., and McCormick, D.A. (2015). Waking State: Rapid Variations Modulate Neural and Behavioral Responses. Neuron 87, 1143–1161. 10.1016/j.neuron.2015.09.012.

55. Reimer, J., Froudarakis, E., Cadwell, C.R., Yatsenko, D., Denfield, G.H., and Tolias, A.S. (2014). Pupil fluctuations track fast switching of cortical states during quiet wakefulness. Neuron 84, 355–362. 10.1016/j.neuron.2014.09.033.

56. Reimer, J., McGinley, M.J., Liu, Y., Rodenkirch, C., Wang, Q., McCormick, D.A., and Tolias, A.S. (2016). Pupil fluctuations track rapid changes in adrenergic and cholinergic activity in cortex. Nat Commun 7, 13289. 10.1038/ncomms13289.

57. Larsen, R.S., and Waters, J. (2018). Neuromodulatory Correlates of Pupil Dilation. Front Neural Circuits 12, 21. 10.3389/fncir.2018.00021.

58. Slater, C., Liu, Y., Weiss, E., Yu, K., and Wang, Q. (2022). The Neuromodulatory Role of the Noradrenergic and Cholinergic Systems and Their Interplay in Cognitive Functions: A Focused Review. Brain Sci 12, 890. 10.3390/brainsci12070890.

59. Gu, Q. (2002). Neuromodulatory transmitter systems in the cortex and their role in cortical plasticity. Neuroscience 111, 815–835. 10.1016/s0306-4522(02)00026-x.

60. Avery, M.C., and Krichmar, J.L. (2017). Neuromodulatory Systems and Their Interactions: A Review of Models, Theories, and Experiments. Front Neural Circuits 11, 108. 10.3389/fncir.2017.00108.

61. Hong, S.Z., Mesik, L., Grossman, C.D., Cohen, J.Y., Lee, B., Severin, D., Lee, H.-K., Hell, J.W., and Kirkwood, A. (2022). Norepinephrine potentiates and serotonin depresses visual cortical responses by transforming eligibility traces. Nat Commun 13, 3202. 10.1038/s41467-022-30827-1.

62. Choi, S.-Y., Chang, J., Jiang, B., Seol, G.-H., Min, S.-S., Han, J.-S., Shin, H.-S., Gallagher, M., and Kirkwood, A. (2005). Multiple receptors coupled to phospholipase C gate long-term depression in visual cortex. J Neurosci 25, 11433–11443. 10.1523/JNEUROSCI.4084-05.2005.

63. He, K., Huertas, M., Hong, S.Z., Tie, X., Hell, J.W., Shouval, H., and Kirkwood, A. (2015). Distinct Eligibility Traces for LTP and LTD in Cortical Synapses. Neuron 88, 528–538. 10.1016/j.neuron.2015.09.037.

64. Bear, M.F., and Singer, W. (1986). Modulation of visual cortical plasticity by acetylcholine and noradrenaline. Nature 320, 172–176. 10.1038/320172a0.

65. Sara, S.J. (2009). The locus coeruleus and noradrenergic modulation of cognition. Nat Rev Neurosci 10, 211–223. 10.1038/nrn2573.

66. Jordan, R., and Keller, G.B. (2023). The locus coeruleus broadcasts prediction errors across the cortex to promote sensorimotor plasticity. eLife 12. 10.7554/eLife.85111.2.

67. Kirkwood, A., Rozas, C., Kirkwood, J., Perez, F., and Bear, M.F. (1999). Modulation of Long-Term Synaptic Depression in Visual Cortex by Acetylcholine and Norepinephrine. J. Neurosci. 19, 1599–1609. 10.1523/JNEUROSCI.19-05-01599.1999.

68. Foote, S.L., and Morrison, J.H. (1987). Extrathalamic modulation of cortical function. Annu Rev Neurosci 10, 67–95. 10.1146/annurev.ne.10.030187.000435.

69. Hurley, L., Devilbiss, D., and Waterhouse, B. (2004). A matter of focus: monoaminergic modulation of stimulus coding in mammalian sensory networks. Current Opinion in Neurobiology 14, 488–495. 10.1016/j.conb.2004.06.007.

70. Kolta, A., Diop, L., and Reader, T.A. (1987). Noradrenergic effects on rat visual cortex: Single-cell microiontophoretic studies of alpha-2 adrenergic receptors. Life Sciences 41, 281–289. 10.1016/0024-3205(87)90150-0.

71. Polack, P.-O., Friedman, J., and Golshani, P. (2013). Cellular mechanisms of brain state– dependent gain modulation in visual cortex. Nat Neurosci 16, 1331–1339. 10.1038/nn.3464.

72. Waterhouse, B.D., Ausim Azizi, S., Burne, R.A., and Woodward, D.J. (1990). Modulation of rat cortical area 17 neuronal responses to moving visual stimuli during norepinephrine and serotonin microiontophoresis. Brain Research 514, 276–292. 10.1016/0006-8993(90)91422-D.

73. Waterhouse, B.D., and Navarra, R.L. (2019). The locus coeruleus-norepinephrine system and sensory signal processing: A historical review and current perspectives. Brain Research 1709, 1–15. 10.1016/j.brainres.2018.08.032.

74. McCormick, D.A. (1989). Cholinergic and noradrenergic modulation of thalamocortical processing. Trends Neurosci 12, 215–221. 10.1016/0166-2236(89)90125-2.

75. Slezak, M., Kandler, S., Veldhoven, P.P.V., Haute, C.V. den, Bonin, V., and Holt, M.G. (2019). Distinct Mechanisms for Visual and Motor-Related Astrocyte Responses in Mouse Visual Cortex. Current Biology 29, 3120–3127.e5. 10.1016/j.cub.2019.07.078.

76. Paukert, M., Agarwal, A., Cha, J., Doze, V.A., Kang, J.U., and Bergles, D.E. (2014). Norepinephrine Controls Astroglial Responsiveness to Local Circuit Activity. Neuron 82, 1263–1270. 10.1016/j.neuron.2014.04.038.

77. Scahill, L. (2009). Alpha-2 adrenergic agonists in children with inattention, hyperactivity and impulsiveness. CNS Drugs 23 Suppl 1, 43–49. 10.2165/00023210-200923000-00006.

78. Franowicz, J.S., Kessler, L.E., Borja, C.M.D., Kobilka, B.K., Limbird, L.E., and Arnsten, A.F.T. (2002). Mutation of the α2A-Adrenoceptor Impairs Working Memory Performance and Annuls Cognitive Enhancement by Guanfacine. J Neurosci 22, 8771–8777. 10.1523/JNEUROSCI.22-19-08771.2002.

79. Posey, D.J., Puntney, J.I., Sasher, T.M., Kem, D.L., and McDougle, C.J. (2004). Guanfacine treatment of hyperactivity and inattention in pervasive developmental disorders: a retrospective analysis of 80 cases. J Child Adolesc Psychopharmacol 14, 233–241. 10.1089/1044546041649084.

80. McKee, J.L., Magielski, J.H., Xian, J., Cohen, S., Toib, J., Chen, C., Kim, D., Rathod, A., Brimble, E., Fitter, N., et al. (2024). Clinical signatures of SYNGAP1-related disorders through data integration. Preprint at medRxiv, 10.1101/2024.10.02.24314452.

81. Kranak, M.P., Rooker, G., and Smith-Hicks, C. (2024). Behavioural phenotype of SYNGAP1-related intellectual disability. Journal of Intellectual Disability Research 68, 1036–1049. 10.1111/jir.13145.

82. Lukkes, J.L., Drozd, H.P., Fitz, S.D., Molosh, A.I., Clapp, D.W., and Shekhar, A. (2020). Guanfacine treatment improves ADHD phenotypes of impulsivity and hyperactivity in a neurofibromatosis type 1 mouse model. J Neurodev Disord 12, 2. 10.1186/s11689-019-9304-y.

83. Artoni, P., Piffer, A., Vinci, V., LeBlanc, J., Nelson, C.A., Hensch, T.K., and Fagiolini, M. (2020). Deep learning of spontaneous arousal fluctuations detects early cholinergic defects across neurodevelopmental mouse models and patients. Proceedings of the National Academy of Sciences 117, 23298–23303. 10.1073/pnas.1820847116.

84. Brain tissue expression of SYNGAP1 - Summary - The Human Protein Atlas https://www.proteinatlas.org/ENSG00000197283-SYNGAP1/brain.

85. Foote, S.L., Bloom, F.E., and Aston-Jones, G. (1983). Nucleus locus ceruleus: new evidence of anatomical and physiological specificity. Physiological Reviews 63, 844–914. 10.1152/physrev.1983.63.3.844.

86. Aston-Jones, G., Ennis, M., Pieribone, V.A., Nickell, W.T., and Shipley, M.T. (1986). The Brain Nucleus Locus Coeruleus: Restricted Afferent Control of a Broad Efferent Network. Science 234, 734–737. 10.1126/science.3775363.

87. Szabadi, E. (2013). Functional neuroanatomy of the central noradrenergic system. J Psychopharmacol 27, 659–693. 10.1177/0269881113490326.

88. Schwarz, L.A., and Luo, L. (2015). Organization of the Locus Coeruleus-Norepinephrine System. Current Biology 25, R1051–R1056. 10.1016/j.cub.2015.09.039.

89. Symposium, U. de M.C. de recherche en sciences neurologiques, Descarries, L., and Reader, T.A. (1984). Monoamine Innervation of Cerebral Cortex: Proceedings of the Fifth Symposium of the Centre de Recherche en Sciences Neurologiques of the Université de Montréal, Held in Montréal, May 16 and 17, 1983 (Liss).

90. Muñoz, W., Tremblay, R., Levenstein, D., and Rudy, B. (2017). Layer-specific modulation of neocortical dendritic inhibition during active wakefulness. Science 355, 954–959. 10.1126/science.aag2599.

91. Toussay, X., Basu, K., Lacoste, B., and Hamel, E. (2013). Locus Coeruleus Stimulation Recruits a Broad Cortical Neuronal Network and Increases Cortical Perfusion. The Journal of Neuroscience 33, 3390. 10.1523/JNEUROSCI.3346-12.2013.

92. Lee, M., Mueller, A., and Moore, T. (2020). Differences in Noradrenaline Receptor Expression Across Different Neuronal Subtypes in Macaque Frontal Eye Field. Front. Neuroanat. 14. 10.3389/fnana.2020.574130.

93. Garcia-Junco-Clemente, P., Tring, E., Ringach, D.L., and Trachtenberg, J.T. (2019). State-Dependent Subnetworks of Parvalbumin-Expressing Interneurons in Neocortex. Cell Reports 26, 2282–2288.e3. 10.1016/j.celrep.2019.02.005.

94. Cardin, J.A., Carlén, M., Meletis, K., Knoblich, U., Zhang, F., Deisseroth, K., Tsai, L.-H., and Moore, C.I. (2009). Driving fast-spiking cells induces gamma rhythm and controls sensory responses. Nature 459, 663–667. 10.1038/nature08002.

95. Sohal, V.S., Zhang, F., Yizhar, O., and Deisseroth, K. (2009). Parvalbumin neurons and gamma rhythms enhance cortical circuit performance. Nature 459, 698–702. 10.1038/nature07991.

96. Carreño-Muñoz, M.I., Chattopadhyaya, B., Agbogba, K., Côté, V., Wang, S., Lévesque, M., Avoli, M., Michaud, J.L., Lippé, S., and Di Cristo, G. (2022). Sensory processing dysregulations as reliable translational biomarkers in SYNGAP1 haploinsufficiency. Brain 145, 754–769. 10.1093/brain/awab329.

97. Chen, G., Zhang, Y., Li, X., Zhao, X., Ye, Q., Lin, Y., Tao, H.W., Rasch, M.J., and Zhang, X. (2017). Distinct Inhibitory Circuits Orchestrate Cortical beta and gamma Band Oscillations. Neuron 96, 1403–1418.e6. 10.1016/j.neuron.2017.11.033.

98. Carlén, M., Meletis, K., Siegle, J.H., Cardin, J.A., Futai, K., Vierling-Claassen, D., Rühlmann, C., Jones, S.R., Deisseroth, K., Sheng, M., et al. (2012). A critical role for NMDA receptors in parvalbumin interneurons for gamma rhythm induction and behavior. Mol Psychiatry 17, 537–548. 10.1038/mp.2011.31.

99. Jones, E.K., Hanley, M., and Riby, D.M. (2020). Distraction, distress and diversity: Exploring the impact of sensory processing differences on learning and school life for pupils with autism spectrum disorders. Research in Autism Spectrum Disorders 72, 101515. 10.1016/j.rasd.2020.101515.

100. Howe, F.E.J., and Stagg, S.D. (2016). How Sensory Experiences Affect Adolescents with an Autistic Spectrum Condition within the Classroom. J Autism Dev Disord 46, 1656–1668. 10.1007/s10803-015-2693-1.

101. Horner, A.E., Norris, R.H., McLaren-Jones, R., Alexander, L., Komiyama, N.H., Grant, S.G.N., Nithianantharajah, J., and Kopanitsa, M.V. (2021). Learning and reaction times in mouse touchscreen tests are differentially impacted by mutations in genes encoding postsynaptic interacting proteins SYNGAP1, NLGN3, DLGAP1, DLGAP2 and SHANK2. Genes, Brain and Behavior 20, e12723. 10.1111/gbb.12723.

102. Vaissiere, T., Michaelson, S.D., Creson, T., Goins, J., Fürth, D., Balazsfi, D., Rojas, C., Golovin, R., Meletis, K., Miller, C.A., et al. (2025). Syngap1 promotes cognitive function through regulation of cortical sensorimotor dynamics. Nat Commun 16, 812. 10.1038/s41467-025-56125-0.

103. Jeyabalan, N., and Clement, J.P. (2016). SYNGAP1: Mind the Gap. Front Cell Neurosci 10, 32. 10.3389/fncel.2016.00032.

104. Ozkan, E.D., Creson, T.K., Kramár, E.A., Rojas, C., Seese, R.R., Babyan, A.H., Shi, Y., Lucero, R., Xu, X., Noebels, J.L., et al. (2014). Reduced cognition in Syngap1 mutants is caused by isolated damage within developing forebrain excitatory neurons. Neuron 82, 1317– 1333. 10.1016/j.neuron.2014.05.015.

105. Clement, J.P., Ozkan, E.D., Aceti, M., Miller, C.A., and Rumbaugh, G. (2013). SYNGAP1 Links the Maturation Rate of Excitatory Synapses to the Duration of Critical-Period Synaptic Plasticity. J. Neurosci. 33, 10447–10452. 10.1523/JNEUROSCI.0765-13.2013.

106. Clement, J.P., Aceti, M., Creson, T.K., Ozkan, E.D., Shi, Y., Reish, N.J., Almonte, A.G., Miller, B.H., Wiltgen, B.J., Miller, C.A., et al. (2012). Pathogenic SYNGAP1 mutations impair cognitive development by disrupting maturation of dendritic spine synapses. Cell 151, 709–723. 10.1016/j.cell.2012.08.045.

107. Kopanitsa, M.V., Gou, G., Afinowi, N.O., Bayés, À., Grant, S.G.N., and Komiyama, N.H. (2018). Chronic treatment with a MEK inhibitor reverses enhanced excitatory field potentials in Syngap1+/− mice. Pharmacological Reports 70, 777–783. 10.1016/j.pharep.2018.02.021.

108. Fenton, T.A., Haouchine, O.Y., Hallam, E.B., Smith, E.M., Jackson, K.C., Rahbarian, D., Canales, C.P., Adhikari, A., Nord, A.S., Ben-Shalom, R., et al. (2024). Hyperexcitability and translational phenotypes in a preclinical mouse model of SYNGAP1-related intellectual disability. Transl Psychiatry 14, 1–11. 10.1038/s41398-024-03077-6.

109. Chandler, D.J. (2015). Evidence for a specialized role of the locus coeruleus noradrenergic system in cortical circuitries and behavioral operations. Brain research 1641, 197. 10.1016/j.brainres.2015.11.022.

110. Agster, K.L., Mejias-Aponte, C.A., Clark, B.D., and Waterhouse, B.D. (2013). Evidence for a regional specificity in the density and distribution of noradrenergic varicosities in rat cortex. J Comp Neurol 521, 2195–2207. 10.1002/cne.23270.

111. Morrison, J.H., Foote, S.L., O’Connor, D., and Bloom, F.E. (1982). Laminar, tangential and regional organization of the noradrenergic innervation of monkey cortex: Dopamine-β-hydroxylase immunohistochemistry. Brain Research Bulletin 9, 309–319. 10.1016/0361-9230(82)90144-7.

112. Kebschull, J.M., Garcia da Silva, P., Reid, A.P., Peikon, I.D., Albeanu, D.F., and Zador, A.M. (2016). High-Throughput Mapping of Single-Neuron Projections by Sequencing of Barcoded RNA. Neuron 91, 975–987. 10.1016/j.neuron.2016.07.036.

113. Guo, X., Hamilton, P.J., Reish, N.J., Sweatt, J.D., Miller, C.A., and Rumbaugh, G. (2009). Reduced expression of the NMDA receptor-interacting protein SynGAP causes behavioral abnormalities that model symptoms of Schizophrenia. Neuropsychopharmacology 34, 1659– 1672. 10.1038/npp.2008.223.

114. Iqbal, E., Govind, R., Romero, A., Dzahini, O., Broadbent, M., Stewart, R., Smith, T., Kim, C.-H., Werbeloff, N., MacCabe, J.H., et al. (2020). The side effect profile of Clozapine in real world data of three large mental health hospitals. PLoS One 15, e0243437. 10.1371/journal.pone.0243437.

115. Berardis, D.D., Rapini, G., Olivieri, L., Nicola, D.D., Tomasetti, C., Valchera, A., Fornaro, M., Fabio, F.D., Perna, G., Nicola, M.D., et al. (2018). Safety of antipsychotics for the treatment of schizophrenia: a focus on the adverse effects of clozapine. Therapeutic Advances in Drug Safety 9, 237. 10.1177/2042098618756261.

116. Prince, J. (2008). Catecholamine dysfunction in attention-deficit/hyperactivity disorder: an update. J Clin Psychopharmacol 28, S39–45. 10.1097/JCP.0b013e318174f92a.

117. Del Campo, N., Chamberlain, S.R., Sahakian, B.J., and Robbins, T.W. (2011). The roles of dopamine and noradrenaline in the pathophysiology and treatment of attention-deficit/hyperactivity disorder. Biol Psychiatry 69, e145–157. 10.1016/j.biopsych.2011.02.036.

118. Wright, D., Kenny, A., Mizen, L.A.M., McKechanie, A.G., and Stanfield, A.C. (2023). Profiling Autism and Attention Deficit Hyperactivity Disorder Traits in Children with SYNGAP1-Related Intellectual Disability. J Autism Dev Disord. 10.1007/s10803-023-06162-9.

119. Nakajima, R., Takao, K., Hattori, S., Shoji, H., Komiyama, N.H., Grant, S.G.N., and Miyakawa, T. (2019). Comprehensive behavioral analysis of heterozygous Syngap1 knockout mice. Neuropsychopharmacology Reports 39, 223–237. 10.1002/npr2.12073.

120. Kilinc, M., Creson, T., Rojas, C., Aceti, M., Ellegood, J., Vaissiere, T., Lerch, J.P., and Rumbaugh, G. (2018). Species-Conserved SYNGAP1 Phenotypes Associated with Neurodevelopmental Disorders. Mol Cell Neurosci 91, 140–150. 10.1016/j.mcn.2018.03.008.

121. Horner, A.E., Norris, R.H., McLaren-Jones, R., Alexander, L., Komiyama, N.H., Grant, S.G.N., Nithianantharajah, J., and Kopanitsa, M.V. (2021). Learning and reaction times in mouse touchscreen tests are differentially impacted by mutations in genes encoding postsynaptic interacting proteins SYNGAP1, NLGN3, DLGAP1, DLGAP2 and SHANK2. Genes, Brain and Behavior 20, e12723. 10.1111/gbb.12723.

122. Samuels, E.R., and Szabadi, E. (2008). Functional Neuroanatomy of the Noradrenergic Locus Coeruleus: Its Roles in the Regulation of Arousal and Autonomic Function Part II: Physiological and Pharmacological Manipulations and Pathological Alterations of Locus Coeruleus Activity in Humans. Current Neuropharmacology 6, 254. 10.2174/157015908785777193.

123. Abercrombie, E.D., and Jacobs, B.L. (1987). Microinjected clonidine inhibits noradrenergic neurons of the locus coeruleus in freely moving cats. Neurosci Lett 76, 203–208. 10.1016/0304-3940(87)90716-6.

124. Aghajanian, G.K., and VanderMaelen, C.P. (1982). α2-Adrenoceptor-Mediated Hyperpolarization of Locus Coeruleus Neurons: Intracellular Studies in Vivo. Science 215, 1394–1396. 10.1126/science.6278591.

125. Marwaha, J., Kehne, J.H., Commissaris, R.L., Lakoski, J., Shaw, W., and Davis, M. (1983). Spinal clonidine inhibits neural firing in locus coeruleus. Brain Research 276, 379–383. 10.1016/0006-8993(83)90752-7.

126. Williams, J.T., Henderson, G., and North, R.A. (1985). Characterization of α2-adrenoceptors which increase potassium conductance in rat locus coeruleus neurones. Neuroscience 14, 95–101. 10.1016/0306-4522(85)90166-6.

127. Jorm, C.M., and Stamford, J.A. (1993). ACTIONS OF THE HYPNOTIC ANAESTHETIC, DEXMEDETOMIDINE, ON NORADRENALINE RELEASE AND CELL FIRING IN RAT LOCUS COERULEUS SLICES. British Journal of Anaesthesia 71, 447–449. 10.1093/bja/71.3.447.

128. Hossmann, V., Maling, T.J.B., Hamilton, C.A., Reid, J.L., and Dollery, C.T. (1980). Sedative and cardiovascular effects of clonidine and nitrazepam. Clinical Pharmacology & Therapeutics 28, 167–176. 10.1038/clpt.1980.146.

129. Hou, R.H., Freeman, C., Langley, R.W., Szabadi, E., and Bradshaw, C.M. (2005). Does modafinil activate the locus coeruleus in man? Comparison of modafinil and clonidine on arousal and autonomic functions in human volunteers. Psychopharmacology 181, 537–549. 10.1007/s00213-005-0013-8.

130. Phillips, M.A., Szabadi, E., and Bradshaw, C.M. (2000). Comparison of the effects of clonidine and yohimbine on spontaneous pupillary fluctuations in healthy human volunteers. Psychopharmacology (Berl) 150, 85–89. 10.1007/s002130000398.

131. Scheinin, M., Kallio, A., Koulu, M., Viikari, J., and Scheinin, H. (1987). Sedative and cardiovascular effects of medetomidine, a novel selective alpha 2-adrenoceptor agonist, in healthy volunteers. British Journal of Clinical Pharmacology 24, 443–451. 10.1111/j.1365-2125.1987.tb03196.x.

132. Nelson, L.E., Lu, J., Guo, T., Saper, C.B., Franks, N.P., and Maze, M. (2003). The α2-Adrenoceptor Agonist Dexmedetomidine Converges on an Endogenous Sleep-promoting Pathway to Exert Its Sedative Effects. Anesthesiology 98, 428–436. 10.1097/00000542-200302000-00024.

133. Heal, D.J., Prow, M.R., and Roger Buckett, W. (1989). Clonidine produces mydriasis in conscious mice by activating central *α*2-adrenoceptors. European Journal of Pharmacology 170, 11–18. 10.1016/0014-2999(89)90127-1.

134. Aoki, C., Go, C.G., Venkatesan, C., and Kurose, H. (1994). Perikaryal and synaptic localization of alpha 2A-adrenergic receptor-like immunoreactivity. Brain Res 650, 181–204. 10.1016/0006-8993(94)91782-5.

135. Sara, S.J., and Bouret, S. (2012). Orienting and Reorienting: The Locus Coeruleus Mediates Cognition through Arousal. Neuron 76, 130–141. 10.1016/j.neuron.2012.09.011.

136. Ji, X.-H., Ji, J.-Z., Zhang, H., and Li, B.-M. (2008). Stimulation of α2-Adrenoceptors Suppresses Excitatory Synaptic Transmission in the Medial Prefrontal Cortex of Rat. Neuropsychopharmacol 33, 2263–2271. 10.1038/sj.npp.1301603.

137. Yi, F., Liu, S.-S., Luo, F., Zhang, X.-H., and Li, B.-M. (2013). Signaling mechanism underlying α2A -adrenergic suppression of excitatory synaptic transmission in the medial prefrontal cortex of rats. Eur J Neurosci 38, 2364–2373. 10.1111/ejn.12257.

138. Nguyen, H.N., Huppé-Gourgues, F., and Vaucher, E. (2015). Activation of the mouse primary visual cortex by medial prefrontal subregion stimulation is not mediated by cholinergic basalo-cortical projections. Frontiers in Systems Neuroscience 9, 1. 10.3389/fnsys.2015.00001.

139. Golmayo, L., Nuñez, A., and Zaborszky, L. (2003). Electrophysiological evidence for the existence of a posterior cortical-prefrontal-basal forebrain circuitry in modulating sensory responses in visual and somatosensory rat cortical areas. Neuroscience 119, 597–609. 10.1016/s0306-4522(03)00031-9.

140. Kang, J.I., Groleau, M., Dotigny, F., Giguère, H., and Vaucher, E. (2014). Visual training paired with electrical stimulation of the basal forebrain improves orientation-selective visual acuity in the rat. Brain Struct Funct 219, 1493–1507. 10.1007/s00429-013-0582-y.

141. Marotta, N., Boland, M.J., and Prosser, B.L. (2024). Accelerating therapeutic development and clinical trial readiness for STXBP1 and SYNGAP1 disorders. Current Problems in Pediatric and Adolescent Health Care 54, 101576. 10.1016/j.cppeds.2024.101576.

142. Aceti, M., Creson, T.K., Vaissiere, T., Rojas, C., Huang, W.-C., Wang, Y.-X., Petralia, R.S., Page, D.T., Miller, C.A., and Rumbaugh, G. (2014). Syngap1 haploinsufficiency damages a postnatal critical period of pyramidal cell structural maturation linked to cortical circuit assembly. Biological psychiatry 77, 805. 10.1016/j.biopsych.2014.08.001.

143. Arora, V., Michaelson, S., Aceti, M., Kilinic, M., Miller, C., and Rumbaugh, G. (2022). Syngap1 Regulates Cortical Circuit Assembly by Controlling Membrane Excitability. Preprint at bioRxiv, 10.1101/2022.12.06.519295.

144. Libé-Philippot, B., Iwata, R., Recupero, A.J., Wierda, K., Garcia, S.B., Hammond, L., Benthem, A. van, Limame, R., Ditkowska, M., Beckers, S., et al. (2024). Synaptic neoteny of human cortical neurons requires species-specific balancing of SRGAP2-SYNGAP1 cross-inhibition. Neuron 112, 3602–3617.e9. 10.1016/j.neuron.2024.08.021.

145. Anderson, J.S., Lodigiani, A.L., Barbaduomo, C.M., and Beegle, J.R. (2024). Hematopoietic stem cell gene therapy for the treatment of SYNGAP1-related non-specific intellectual disability. The Journal of Gene Medicine 26, e3717. 10.1002/jgm.3717.

146. Porter, K., Komiyama, N.H., Vitalis, T., Kind, P.C., and Grant, S.G.N. (2005). Differential expression of two NMDA receptor interacting proteins, PSD-95 and SynGAP during mouse development. European Journal of Neuroscience 21, 351–362. 10.1111/j.1460-9568.2005.03874.x.

147. Komiyama, N.H., Watabe, A.M., Carlisle, H.J., Porter, K., Charlesworth, P., Monti, J., Strathdee, D.J.C., O’Carroll, C.M., Martin, S.J., Morris, R.G.M., et al. (2002). SynGAP regulates ERK/MAPK signaling, synaptic plasticity, and learning in the complex with postsynaptic density 95 and NMDA receptor. J Neurosci 22, 9721–9732. 10.1523/JNEUROSCI.22-22-09721.2002.

148. Oliveira, L.S., Sumera, A., and Booker, S.A. (2021). Repeated whole-cell patch-clamp recording from CA1 pyramidal cells in rodent hippocampal slices followed by axon initial segment labeling. STAR Protoc 2, 100336. 10.1016/j.xpro.2021.100336.

149. Booker, S.A., Song, J., and Vida, I. (2014). Whole-cell Patch-clamp Recordings from Morphologically- and Neurochemically-identified Hippocampal Interneurons. J Vis Exp, 51706. 10.3791/51706.

150. Pakan, J.M., Lowe, S.C., Dylda, E., Keemink, S.W., Currie, S.P., Coutts, C.A., and Rochefort, N.L. Behavioral-state modulation of inhibition is context-dependent and cell type specific in mouse visual cortex. eLife 5, e14985. 10.7554/eLife.14985.

151. Wong, A.A., and Brown, R.E. (2006). Visual detection, pattern discrimination and visual acuity in 14 strains of mice. Genes, Brain and Behavior 5, 389–403. 10.1111/j.1601-183X.2005.00173.x.

152. Guzman, S.J., Schlögl, A., and Schmidt-Hieber, C. (2014). Stimfit: quantifying electrophysiological data with Python. Front Neuroinform 8, 16. 10.3389/fninf.2014.00016.

153. Longair, M.H., Baker, D.A., and Armstrong, J.D. (2011). Simple Neurite Tracer: open source software for reconstruction, visualization and analysis of neuronal processes. Bioinformatics 27, 2453–2454. 10.1093/bioinformatics/btr390.

154. Henschke, J.U., Dylda, E., Katsanevaki, D., Dupuy, N., Currie, S.P., Amvrosiadis, T., Pakan, J.M.P., and Rochefort, N.L. (2020). Reward Association Enhances Stimulus-Specific Representations in Primary Visual Cortex. Current Biology 30, 1866–1880.e5. 10.1016/j.cub.2020.03.018.

155. Kaifosh, P., Zaremba, J.D., Danielson, N.B., and Losonczy, A. (2014). SIMA: Python software for analysis of dynamic fluorescence imaging data. Front Neuroinform 8, 80. 10.3389/fninf.2014.00080.

156. Schindelin, J., Arganda-Carreras, I., Frise, E., Kaynig, V., Longair, M., Pietzsch, T., Preibisch, S., Rueden, C., Saalfeld, S., Schmid, B., et al. (2012). Fiji: an open-source platform for biological-image analysis. Nat Methods 9, 676–682. 10.1038/nmeth.2019.

157. Keemink, S.W., Lowe, S.C., Pakan, J.M.P., Dylda, E., van Rossum, M.C.W., and Rochefort, N.L. (2018). FISSA: A neuropil decontamination toolbox for calcium imaging signals. Sci Rep 8, 3493. 10.1038/s41598-018-21640-2.

158. Mazurek, M., Kager, M., and Van Hooser, S.D. (2014). Robust quantification of orientation selectivity and direction selectivity. Front Neural Circuits 8, 92. 10.3389/fncir.2014.00092.

159. Alexander, R.A. (1990). A note on averaging correlations. Bull. Psychon. Soc. 28, 335–336. 10.3758/BF03334037.

160. Corey, D.M., Dunlap, W.P., and Burke, M.J. (1998). Averaging Correlations: Expected Values and Bias in Combined Pearson rs and Fisher’s z Transformations. The Journal of General Psychology 125, 245–261. 10.1080/00221309809595548.

161. Montijn, J.S., Vinck, M., and Pennartz, C.M.A. (2014). Population coding in mouse visual cortex: response reliability and dissociability of stimulus tuning and noise correlation. Frontiers in Computational Neuroscience 8.

162. Dadarlat, M.C., and Stryker, M.P. (2017). Locomotion Enhances Neural Encoding of Visual Stimuli in Mouse V1. J. Neurosci. 37, 3764–3775. 10.1523/JNEUROSCI.2728-16.2017.

163. Mathis, A., Mamidanna, P., Cury, K.M., Abe, T., Murthy, V.N., Mathis, M.W., and Bethge, M. (2018). DeepLabCut: markerless pose estimation of user-defined body parts with deep learning. Nat Neurosci 21, 1281–1289. 10.1038/s41593-018-0209-y.

164. Fan, X., Miles, J.H., Takahashi, N., and Yao, G. (2009). Abnormal Transient Pupillary Light Reflex in Individuals with Autism Spectrum Disorders. J Autism Dev Disord 39, 1499–1508. 10.1007/s10803-009-0767-7.

165. Nyström, P., Gredebäck, G., Bölte, S., Falck-Ytter, T., and EASE team (2015). Hypersensitive pupillary light reflex in infants at risk for autism. Molecular Autism 6, 10. 10.1186/s13229-015-0011-6.

